# Modeling Post-Gastrula Development via Bidirectional Pluripotent Stem Cells

**DOI:** 10.1101/2025.06.28.662107

**Authors:** Kuisheng Liu, Zihui Yan, Dandan Bai, Rui Jiang, Yan Bi, Xiangjun Ma, Jiani Xiang, Yifan Sheng, Baoxing Dong, Zhiyuan Ning, Shanru Yi, Yingdong Liu, Xinyi Lei, Yanping Jia, Yan Zhang, Yalin Zhang, Yanhe Li, Chenxiang Xi, Shanyao Liu, Shuyi Liu, Jiayu Chen, Jiqing Yin, Xiaochen Kou, Yanhong Zhao, Hong Wang, Yixuan Wang, Ke Wei, Wenqiang Liu, Shaorong Gao

## Abstract

The absence of stem cells capable of efficiently generating both trophoblast and epiblast lineages has hindered precise recapitulation of embryonic development. Through high-content chemical screening, we established an AL medium to generate mouse and human bidirectional pluripotent stem cells (BPSCs) characterized by concurrent OCT4/CDX2 expression. Mouse BPSCs demonstrated high plastic differentiation into trophoblast, epiblast and primitive endoderm lineages in vitro within 48 hours without exogenous induction factors and efficiently contributed to the embryo and extraembryonic tissues in vivo. Mechanistically, hyperactivation of the Wnt signaling pathway breaks the early lineage differentiation barrier by initiating a Lef1-dependent bypass. Remarkably, BPSCs can efficiently generated E8.5 embryoids that completed gastrulation and displayed advanced features such as brain development, a closed neural tube, a beating heart, somite formation, and primordial germ cells. These findings highlight BPSCs as a powerful tool for investigating early lineage specification and post-gastrulation embryonic development, with potential applications across multiple species.

## Introduction

Mammalian embryos undergo two critical cell fate determinations, resulting in the formation of three preimplantation lineages: trophectoderm (TE), epiblast (Epi), and primitive endoderm (PrE)^1–4^. This process is foundational for embryo implantation, germ layer differentiation, and yolk sac and placenta formation^5,6^. However, studying early embryogenesis is challenging owing to the limited availability of embryos and the complex nature of embryo implantation. Previous studies have successfully derived embryonic stem cells (ESCs), trophoblast stem cells (TSCs), and extraembryonic endoderm stem (XEN) cell lines that represent early lineages, thereby facilitating the study of embryonic development^7–10^. Totipotent-like stem cells that can bidirectional develop into both trophoblast and epiblast lineages offer a promising tool for modeling embryogenesis.

Due to the establishment of lineage barriers, the conversion of ESCs to TSCs often necessitates the forced expression of exogenous transcription factors (CDX2)^11,12^. Although recent studies have reported that changing the culture conditions can convert totipotent-like cells or ESCs into TSCs, the process remains lengthy and inefficient^13–23^. Currently, during the process of acquiring the developmental potential of the trophoblast lineage, pluripotent cells often have to endure significant epigenetic perturbations or cellular stress, which can lead to cell death or slowed proliferation. Therefore, breaking the developmental barriers between cell lineages and achieving efficient interconversion among ESCs, TSCs, and XEN cell lines could contribute to a better understanding of lineage-determining processes during embryonic development.

The construction of an embryo-like structure is a method for recapitulating embryonic developmental processes through 3D co-culture of multiple cell lines. Currently, it is possible to produce blastoids (blastocyst-like structures), gastrulating embryo-like structures (embryonic day 6.5-like structure, E6.5-like structure), and post-gastrulation synthetic embryos (E8.5-like structure) by co-culturing multiple cells^24–33^. However, the method of using a mixture of different cell lines to construct post-implantation embryo-like structures is complex and inefficient, and there is also the risk of inconsistent genetic backgrounds, which seriously affects its further application. Thus, constructing embryo-like structures simply and efficiently from a single cell line would greatly advance this field.

Here, through high-content screening, we identified a cell culture condition that could easily and nondestructively enable mouse and human pluripotent stem cells to express both *Oct4* and *Cdx2*, and designated these cells as bidirectional pluripotent stem cells (BPSCs). Further studies have revealed that BPSCs have bidirectional developmental potential for efficiently contributing to both the embryonic and extraembryonic lineages. In addition, the conversion of BPSCs into TSCs was achieved efficiently. During the construction of embryo-like structures, the BPSCs demonstrated strong constructive ability and successfully achieved E5.5–E8.5 like embryos. This study broadens our understanding of potential bidirectional cell states and further provides a simple, robust and highly efficient embryoid model for mimicking post-implantation embryogenesis.

## Results

### Chemical induction of bidirectional pluripotent stem cells (BPSCs) with double-positive of OCT4 and CDX2

To screen for culture conditions that facilitate the differentiation of pluripotent stem cells into trophoblast lineage cells, we first established extended pluripotent stem cell (EPSC) lines with *Oct4*-GFP and CDX2-mCherry fluorescent reporters. After culturing the EPSCs in N2B27 basal medium supplemented with chemical libraries for 3 days, the intensity of *Oct4*-GFP and CDX2-mCherry fluorescence was detected (Figure 1A). Out of the 1,859 small molecules tested, 261 enhanced *Oct4*-GFP fluorescence compared to the control, but only 10 increased CDX2-mCherry fluorescence (Figure 1B; Table S1). To our surprise, there are two small molecules, LY2090314 (LY) and AS1842856 (AS), could promote the expression of OCT4 and CDX2 simultaneously in a concentration-dependent manner (Figures 1B, S1A and S1B). After further experiments, a high proportion of *Oct4*-GFP and CDX2-mCherry double-positive cells was induced by AS at 0.6 μM (66.03%) and LY at 10 nM (53.67%) with low cell-toxicity (Figure 1C). When cotreated with AS and LY (AL) for 3 days, the proportion of CDX2 and OCT4 double-positive cells increased to 70.65% (Figure 1C). ESCs also exhibited a significant response to AL treatment, with more than 60% enrichment of CDX2 and OCT4 double-positive cells (Figure 1C). Notably, AL-treated EPSCs produced dome-shaped colonies similar to those of ESCs and EPSCs (Figure 1D). The results of immunofluorescence showed that OCT4 and CDX2 were colocalized in the nucleus of double-positive cells, and there was no obvious mutual exclusion phenomenon (Figure 1E). Nuclear staining and flow cytometry showed that the proportion of cells co-expressing OCT4 and CDX2 proteins exceeded 70% (Figure 1F).

**Figure 1.**
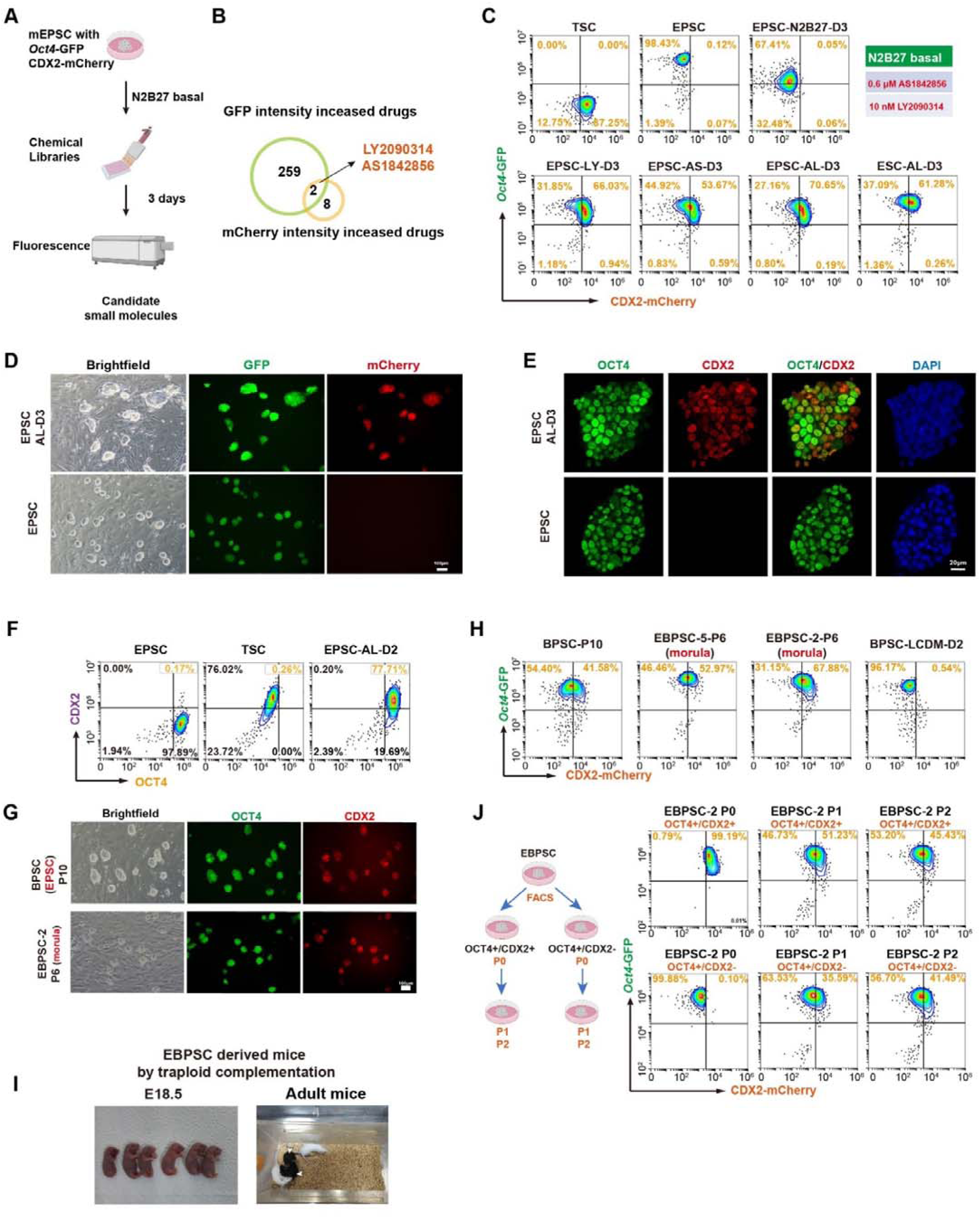
Chemical induction of bidirectional pluripotent stem cells (BPSCs) with double-positive of OCT4 and CDX2. (A) Schematic representation of chemical library screening for *Oct4*-GFP and CDX2-mCherry double-positive cells. (B) The Venn diagram shows that 261 small molecules significantly increased GFP expression and 10 significantly increased mCherry expression compared to the negative control. Two small molecules, LY2090314 and AS1842856, were common to both groups. (C) Representative FACS analysis of the percentages of *Oct4*-GFP positive and CDX2-mCherry positive cells in TSC, EPSC, EPSC with differentiated in N2B27 medium for 3 days (EPSC-N2B27-D3), EPSC treated with LY for 3 days (EPSC-LY-D3), EPSC treated with AS for 3 days (EPSC-AS-D3) and EPSC co-treated with both AS and LY (AL) for 3 days (EPSC-AL-D3), ESC co-treated with both AS and LY (AL) for 3 days (ESC-AL-D3). (D) Representative morphological images and *Oct4*-GFP and CDX2-mCherry fluorescence in EPSC-AL-D3 and EPSC. Scale bar, 100 μm. (E) Immunofluorescence staining of EPSC and EPSC-AL-D3. Staining for CDX2 and OCT4. Scale bar, 20 μm. (F) Representative FACS analysis of nuclear proteins OCT4 and CDX2 of EPSC, TSC and EPSC-AL-D2. (G) Representative morphological images and *Oct4*-GFP, CDX2-mCherry fluorescence of BPSC (converted from EPSCs and cultured for 10 passages) and EBPSC (derived from morula and cultured for 6 passages). Scale bar, 100 μm. (H) Representative FACS analysis of the percentages of *Oct4*-GFP positive and CDX2-mCherry positive cells in BPSC derived from EPSC and cultured for 10 passages (BPSC-P10), BPSC derived from morula (16- to 32-cell) stage and cultured for 6 passages (EBPSC-5-P6, EBPSC-2-P6), and BPSC cultured with EPSC medium (LCDM) for 2 days (BPSC-LCDM-D2). (I) Brightfield image of tetraploid complemented embryos at day E18.5 and adult mice. (J) Schematic diagram (left panel) showing BPSC flow sorting and culture for two passages. Representative FACS analysis of the percentages of Oct4-GFP positive and CDX2-mCherry positive cells (right panel).

Mouse embryos also exhibit co-expression of key transcription factors prior to the first cell fate determination^11,34–36^. The single-cell sequencing data and protein detection on mouse preimplantation embryonic development revealed that the critical transcription factors *Cdx2*, and *Oct4* were expressed at the 8-cell stage and highly expressed at the 16-cell stage (Figures S1C and S1D). During the E3.5 blastocyst stage, these transcription factors were co-expressed in TE, and they will be completely separated at late blastocyst stage (E4.5) (Figure S1D). Previous studies have demonstrated that the outer blastomeres of the morula can contribute to both embryonic and extra-embryonic lineages, indicating that cells showing co-localization of CDX2 and OCT4 possess bidirectional developmental potential^37^. We then designated these AL-treated cells as bidirectional pluripotent stem cells (BPSCs).

We next examined whether the BPSCs could be maintained in vitro for a long period of time. Culture and fluorescence-activated cell sorting (FACS) analyses showed that BPSCs retained up to 41.58% of CDX2 and OCT4 double-positive cells and maintained dome-shaped colonies over 10 passages (Figures 1G and 1H). Moreover, we derived BPSCs from embryos at the morula (16- to 32-cell) stage. Two of three cell lines (EBPSC-2 and EBPSC-5) showed a normal karyotype over 10 passages and a high percentage (> 50%) of cells double-positive for OCT4 and CDX2 over six passages (Figures 1G, 1H and S1E). To verify the developmental potential of BPSCs, we performed tetraploid complementation experiments. The results showed that EBPSCs could generate tetraploid-complemented mice and develop normally to adulthood (Figures 1I and S1F). These results indicated that this culture system is robust for the derivation and long-term maintenance of BPSCs with whole embryo development potential.

To interrogate the stability of the proportion of CDX2 and OCT4 double-positive cell composition, we performed flow sorting single-positive (Oct4+/Cdx2-) and double-positive (Oct4+/Cdx2+) cells subpopulations from BPSCs and maintained them separately in AL medium (P0). Strikingly, by passage 1 (P1), both subpopulations re-established the original equilibrium of double-positive cells (Figure 1J). In addition, when BPSCs derived from EPSCs were re-cultured in EPSCs medium LCDM, most of the cells (96%) returned to the single-positive state (Figure 1H). This indicates that BPSCs maintain a highly plastic state under AL culture conditions, allowing for dynamic transitions between transcriptional states.

### BPSCs possess high potential for embryonic and extraembryonic lineages

Previous studies have shown that mouse pluripotent stem cells or totipotent-like stem cells cannot rapidly transform into TSCs without ectopic expression of the TE core transcription factor CDX2^11–15,18,19,21–23,38^. To investigate the developmental potential of BPSCs *in vitro*, we cultured BPSCs, ESCs and EPSCs under inducing-factors-free serum medium conditions (Figure 2A). After 2 days, single cells positive for CDX2, OCT4, or GATA6 were detected in the autonomously differentiating clones (Figures 2B and 2C). In contrast to the low efficiency observed for transformed TSCs in EPSCs and ESCs, BPSCs markedly increased the yield of CDX2-expressing cells to 40.36% (Figure 2D).

**Figure 2.**
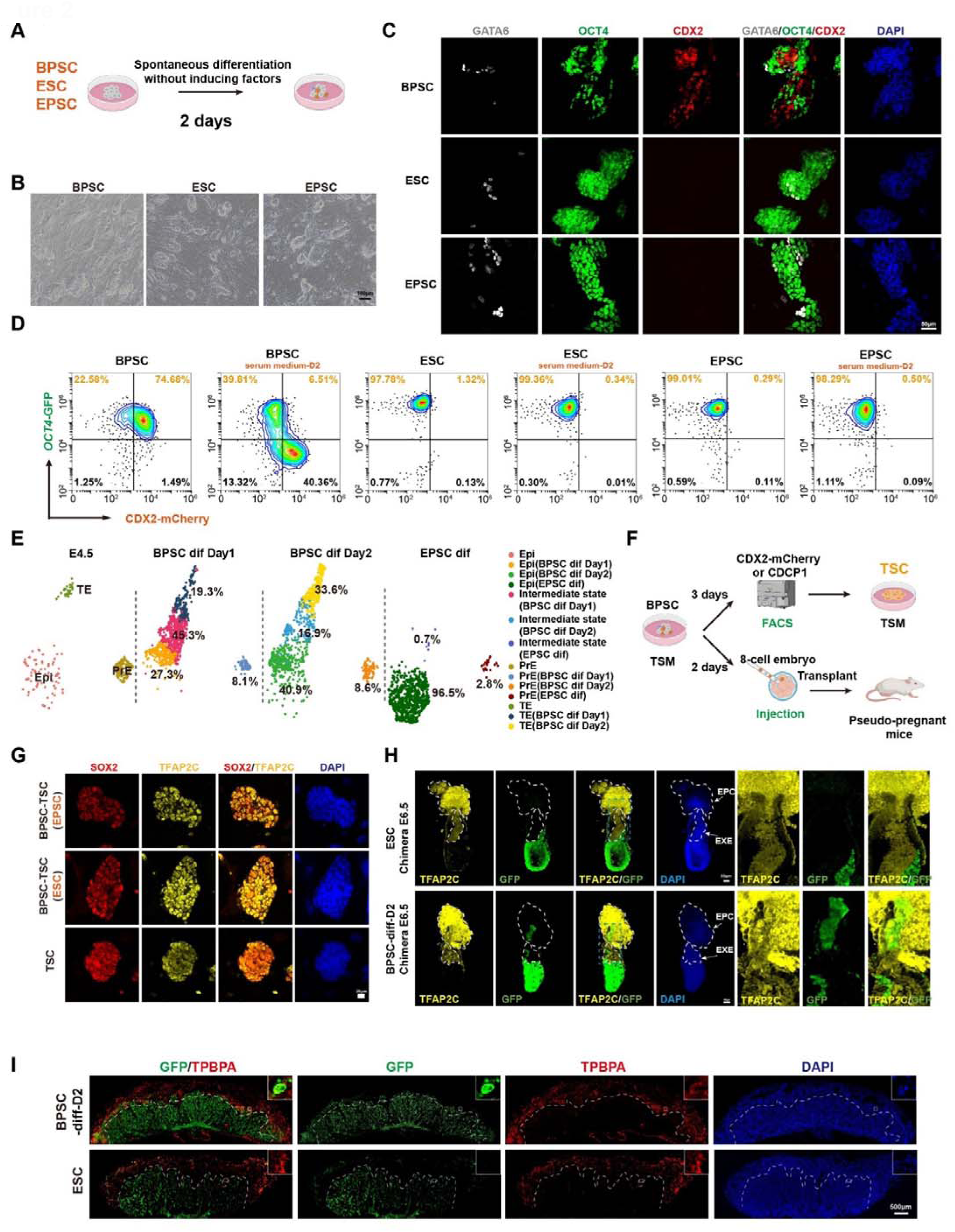
BPSCs possess high potential for embryonic and extraembryonic lineages. (A) Schematic of BPSC (derived from EPSC), ESC, and EPSC differentiated spontaneously without inducing factors for 2 days. (B) Typical brightfield images of BPSC (derived from EPSC), ESC, and EPSC after two days of spontaneous differentiation without inducing factors. (C) Immunofluorescence staining of BPSC (derived from EPSC), ESC, and EPSC differentiated spontaneously for 2 days. Staining for OCT4, CDX2 and GATA6. Scale bar, 50 μm. (D) Representative FACS analysis of the BPSC (derived from EPSC), ESC, and EPSC differentiated spontaneously for 2 days. (E) UMAP plot of scRNA-seq data from E4.5 embryo cells, BPSC (derived from EPSC) after 1 or 2 days of spontaneous differentiation, and EPSC after 2 days of spontaneous differentiation. Cells were colored according their cell types and split by their origins. (F) Schematic of BPSC differentiation induced with TSM for 2-3 days. After 3 days of induction, CDX2-mCherry-positive or CDCP1-positive cells were collected by FACS and cultured with TSM. After 2 days of induction, chimeras were performed with cells injected into embryos at 8-cell stage. (G) Immunofluorescence staining of BPSC-TSC (derived from EPSC), BPSC-TSC (derived from ESC) and TSCs (derived from embryos). Staining for SOX2 and TFAP2C. Scale bar, 20 μm. (H) Immunofluorescence staining of E6.5 chimeric embryo produced using the above method (Figure 3F). Staining for TFAP2C and GFP. Scale bar, 50 μm. BPSC and ESC are labeled by GFP. Enlarged view of the blue dotted box (right panel). (I) Immunofluorescence staining of chimeric placentas at E12.5 stage. Staining for TPBPA and GFP. Scale bar, 500 μm.

To explore the developmental status, we performed UMAP analysis of scRNA-seq data from autonomously differentiating cells of BPSCs and EPSCs (Figure 2E). Compared to E4.5 blastocyst^39^, the data revealed that TE- (33.6%), PrE- (8.6%), and Epi-like cells (40.9%) in autonomously differentiating cells (day2) of BPSCs mapped separately to the TE, PrE, and Epi of the E4.5 blastocysts, with robust expression of lineage markers (Figures 2E and S2A). Unlike the previously reported totipotent like stem cells, more than 50% of BPSCs immediately differentiate into the Epi-, TE-, and PrE-like lineages on the first day of differentiation. In contrast, EPSCs only involved Epi- (96.5%) and PrE-like cells (2.8%), but not TE-like cells (Figure 2E).

As BPSCs consist of both single-positive (*Oct4*+/*Cdx2*-) and double-positive (*Oct4*+/*Cdx2*+) cells, it remains unclear whether these two cell types differ in their differentiation directions. We sorted out the two types of cells in BPSCs and discarded the PrE lineage cells using PDGFRA antibodies (Figure S2B). The two groups of cells were subjected to autonomous differentiation experiments, and the results showed that double-positive cells were the main source of TE-like lineage cells, while single-positive cells rarely differentiated into TE-like lineage cells, and PrE-like lineage cells were mostly differentiated from OCT4 single-positive cells (Figure S2C). In addition, we performed single-cell differentiation of flow-sorted cells. Unlike their autonomous differentiation capacity in bulk culture, double-positive cells require inducing factors (FGF4 and Heparin) to differentiate into TE-like cells under single-cell conditions (Figures S2D and S2E). This phenomenon may be due to the lack of intercellular communication in single-cell differentiation, which is present in bulk-cell differentiation and can determine the fate of individual cells as they differentiate autonomously. Interestingly, most of these clones contain only one cell type under single-cell differentiation condition, suggesting that BPSC immediately determines the direction of differentiation in the case of single cells (Figures S2D and S2E).

To determine whether TSC lines could be generated, BPSCs were treated with a directed induction medium (TSM with Fgf4 and Heparin) for 3 days and detected with CDX2-mCherry or CDCP1 (a surface marker of TSCs) (Figure 2F). Similar to EPSC-derived BPSCs (EPSC-BPSC-TSMD3), ESC-derived BPSCs (ESC-BPSC-TSMD3) responded to TSM induction and produced up to 60.55% CDCP1-positive cells (Figure S3A). After collection by FACS (CDX2-mCherry-positive or CDCP1-positive cells) and culture with TSM for 3–4 days, the TSC lines were successfully established. Compared with embryo-derived TSCs, the BPSC-converted TSC lines (BPSC-TSC(EPSC) and BPSC-TSC(ESC)) showed tight epithelial clones and expressed the TSC markers SOX2, TFAP2C, and CDX2 (Figures 2G and S3B).

To explore the differentiation potential of BPSCs *in vivo*, differentiated cells treated with TSM for 2 days (labeled with GFP) were injected into 8-cell stage embryos (Figure 2F). Chimeras were examined at E6.5, and 30–45% of the embryos displayed a clear contribution of BPSCs (GFP-positive) in Epi, extraembryonic ectoderm (ExE), and ectoplacental cone (EPC) (Figures 2H, S3C and S3D). In contrast, only Epi exhibited GFP fluorescence signals in chimeras generated using ESCs (Figures 2H and S3C). Additionally, a stable bidirectional contribution of BPSCs to fetal and extraembryonic tissues (placenta and yolk sac) was detected in chimeras at the E12.5 stage (Figures 2I, S3E).

These findings indicate that BPSCs possess extraordinary plasticity in terms of bidirectional differentiation potential for the development of embryonic and extraembryonic lineages.

### BPSCs exhibit a unique pluripotency state

To investigate the composition of BPSCs, we collected scRNA-seq data and performed uniform manifold approximation and projection (UMAP) analysis with EPSCs and ESCs (Figures 3A, 3B and S4A). On a global transcriptomic level, BPSCs differ from both EPSCs and ESCs and include a small fraction of PrE-like cells. BPSCs highly express a subset of TE lineage genes, such as *Cdx2*, *Krt8*, *Krt18*, while pluripotency genes remain largely unchanged (Figure 3B and S4A). Notably, BPSCs contain a substantial population of cells that co-express *Oct4* and *Cdx2* (Figure 3B). To elucidate the basis for the high efficiency of TE lineage differentiation in BPSCs, we assessed the genomic accessibility and profiled histone modifications, including H3K27ac, H3K4me3, and H3K27me3, in BPSCs, followed by comparative analysis (Figures S4B-D). Comparing BPSCs with EPSCs and ESCs, we found that although BPSCs do not broadly express a large number of TE lineage genes at the transcriptomic level, these genes exhibit greater chromatin accessibility in BPSCs (Figure 3C, 3D and 3E). This pre-opened chromatin state corresponds to the rapid differentiation towards extraembryonic lineages. Furthermore, analysis of motif accessibility for key TE- and ICM-lineage transcription factors revealed increased accessibility in BPSCs for genomic regions containing CDX2, TFAP2C, EOMES, and TEAD4 motifs, while accessibility for NANOG and OCT4 motifs was unchanged (Figure 3F). Regarding histone modifications, there is a notable upregulation of H3K27ac at the TE lineage markers in BPSCs, while H3K4me3 and H3K27me3 exhibited a slight downregulation (Figure S4E).

**Figure 3.**
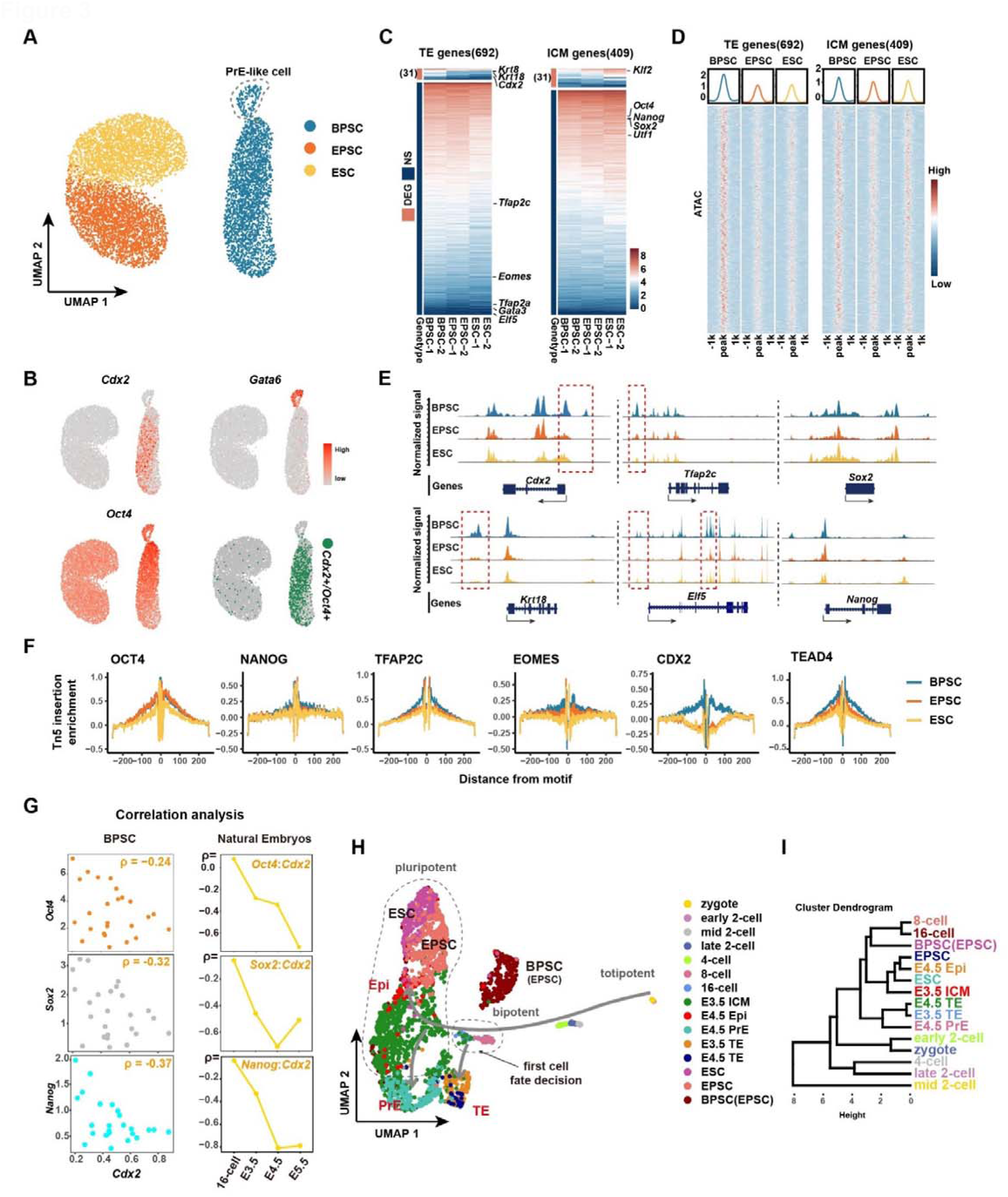
BPSCs exhibit a unique pluripotency state. (A) UMAP plot of scRNA-seq data from BPSC, EPSC, and ESC. (B) Gene expression of *Cdx2*, *Gata6*, and *Oct4* in BPSC, EPSC and ESC. Cells identified as *Cdx2*+/*Oct4*+ are also labeled (bottom right). (C) Heatmap showing the expression of TE and ICM marker genes in BPSC, EPSC, and ESC. The expression of genes is normalized by TPM and log-transformed. Genes identified as differentially expressed genes (DEG) or non-significant genes (NS) between BPSCs and either ESC or EPSC, the representative genes are shown on the left side of the heatmap. (D) Chromatin accessibility signal heatmap centered on peaks (±1 kb) in the promoter region of TE and ICM marker genes in BPSC, EPSC, and ESC. Each row corresponds to an individual peak, and color scale indicates normalized accessibility signal. Line plots above the heatmap show the average accessibility profiles around peaks (E) Genome browser view of chromatin accessibility signal distribution on the region of representative TE and ICM marker genes. Red dotted line indicates the differential chromatin accessibility signal between BPSC and EPSC/ESC. (F) Transcription factor footprint analysis based on scATAC-seq data of BPSC, EPSC, and ESC. The plot shows the normalized Tn5 insertion frequency centered on the motif (±250 bp) of several transcription factors. Different lines represent different cell types. (G) Scatter plot showing the expression levels between *Cdx2* and *Oct4*, *Sox2*, *Nanog*, respectively in BPSCs. Each dot represents one cluster in BPSC divided according to the scRNA-seq and the gene expression values are calculated as average expressions. The Spearman’s rank correlation coefficients (ρ) are also listed. The line chart shows the Spearman’s rank correlation coefficients between *Cdx2* and *Oct4*, *Sox2*, *Nanog*, respectively in different stages of natural embryos. (H) UMAP plot of scRNA-seq data from BPSC, EPSC and ESC, and pre-implantation mouse embryos from zygote to E4.5 blastocyst. (I) Hierarchical clustering analysis of BPSC, EPSC, ESC, and pre-implantation mouse embryos from zygote to E4.5 blastocyst.

Similar to the FACS and immunofluorescent staining results, Single-cell transcriptomic analysis revealed that approximately 48% of the cells co-expressed *Oct4* and *Cdx2* and the remaining cells are primarily OCT4 single-positive cells (Figures S4F and S4G). BPSCs were homogeneous with scattered expression of the trophoblast-related gene, *Cdx2*, except for a few PrE-like cells (Figure 3B). Despite the typical mutual exclusivity of *Oct4* and *Cdx2* in pluripotent stem cells, their correlation in BPSCs and preimplantation embryos remains unexplored. We found a negative correlation between *Oct4* and *Cdx2* from E3.5 to E5.5, but not at the 16-cell stage (Figure 3G). BPSCs showed a similar weak negative correlation between these two genes as observed in 16-cell embryos (Figure 3G). Moreover, the correlations between *Sox2:Cdx2* and *Nanog:Cdx2* in BPSCs were more like those in morula (16- to 32-cell) stage embryos than those after E3.5 (Figure 3G).

To compare BPSCs with preimplantation embryos, we analyzed single cell RNA-seq data with published preimplantation embryo data. UMAP showed that BPSCs were similar to embryos at the morula stage (16-cell) which marks the onset of the first cell fate determination period (Figures 3H and S4H). In contrast, ESCs and EPSCs are closer to ICM (E3.5) and Epi (E4.5) (Figure 3H). Further clustering analysis showed that BPSCs clustered with 8- and 16-cell embryos (Figure 3I).

### Hyperactivation of Wnt signalling pathway expands the developmental boundaries of pluripotent stem cells by activate *Lef1*

To investigated the mechanism of AL medium confers extraembryonic confers extraembryonic differentiation competence, transcriptomic profiling revealed significant upregulation of Wnt pathway components following AS, LY, or combined treatment (Figure 4A). Additionally, we tested another canonical Wnt activator, CHIR. Compared to a low concentration (3 μM) used for maintaining pluripotency, only higher concentration (10 μM) of CHIR activated CDX2 expression (Figures S5A and S5B), suggestion dose-dependent Wnt activation as critical for lineage conversion.

**Figure 4.**
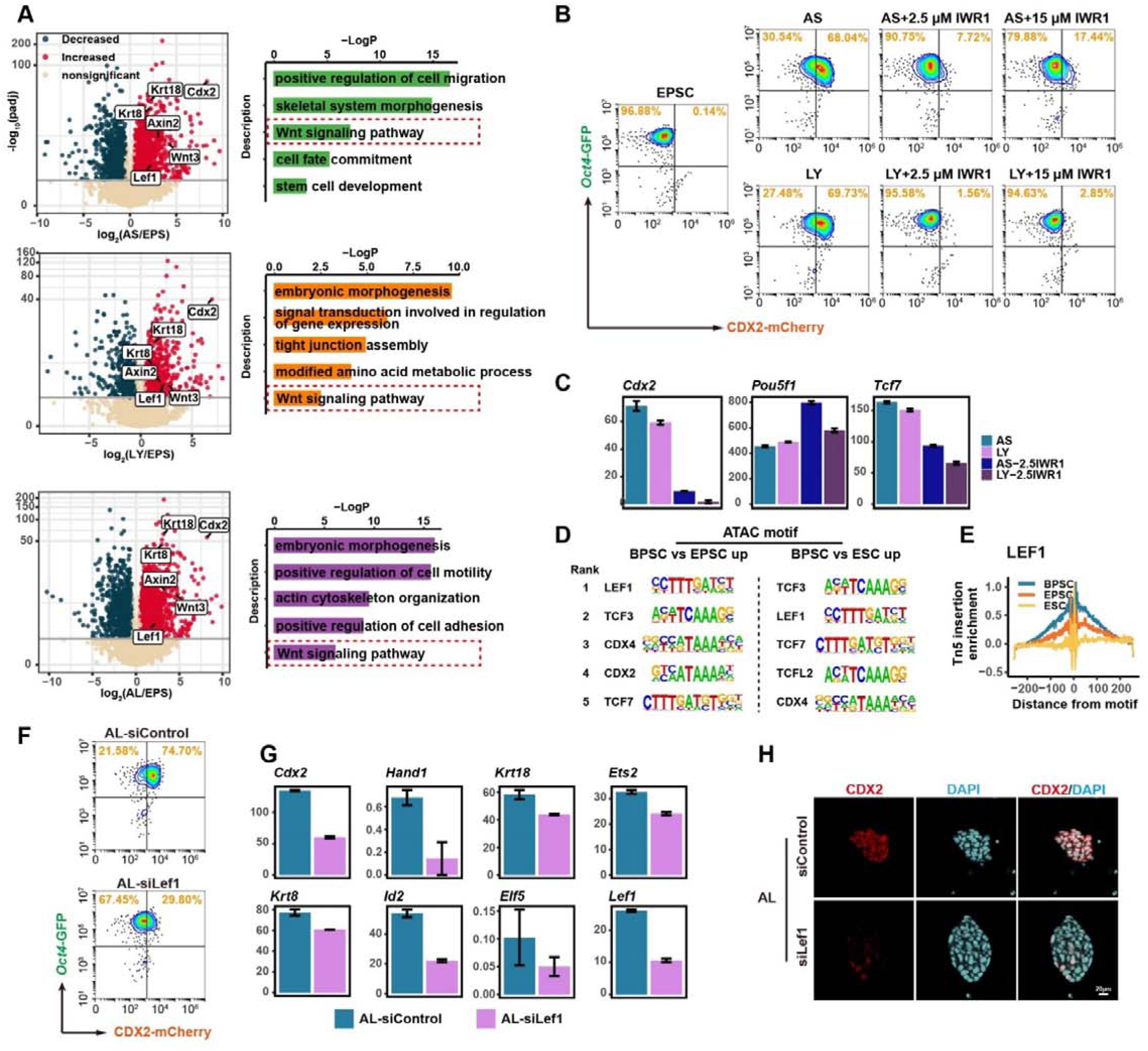
Wnt signaling pathway expands the developmental boundaries of pluripotent stem cells. (A) Volcano plot showing the differentially expressed genes between EPSC treated with AS, LY, or AS+LY (AL) and untreated EPSC (left panel), and the GO enrichment analysis of upregulated genes (right panel). (B) Representative FACS analysis of the percentages of Oct4-GFP positive and CDX2-mCherry positive cells in EPSC (LCDM), AS, AS+2.5 μM IWR1, AS+15 μM IWR1, LY, LY+2.5 μM IWR1, LY+15 μM IWR1. (C) Bar plot showing the expression of *Cdx2*, *Oct4*, and *Tcf7* in the bulk RNA-seq data of EPSC under AS, LY, AS+2.5 μM IWR1, LY+2.5 μM IWR1 condition. The expression of genes is normalized by TPM. (D) Top 5 enriched transcription factors identified based on motif analysis from upregulated scATAC-seq peaks in BPSC compared with EPSC or ESC. (E) Transcription factor footprint analysis of *Lef1* based on scATAC-seq data of BPSC, EPSC, and ESC. The plot shows the normalized Tn5 insertion frequency centered on the motif (±250 bp) of several transcription factors. Different lines represent different cell types. (F) Representative FACS analysis of the percentages of Oct4-GFP positive and CDX2-mCherry positive cells in BPSC (AL) under knockdown control and knockdown *Lef1* condition. (G) Bar plot showing the expression of several TE marker genes in the bulk RNA-seq data of *Lef1* knockdown group and control group of AL treated EPSC. The expression of genes is normalized by TPM. (H) Immunofluorescence staining of BPSC (AL) under knockdown control and knockdown Lef1 condition. Staining for CDX2. Scale bar, 20 μm.

To further elucidate the role of Wnt signaling pathway, we added the Wnt inhibitor IWR1 to N2B27 basal medium containing AS and LY. Flow cytometry and transcriptomic analyses showed that even low concentrations of IWR1 significantly inhibited AS- and LY-induced CDX2 activation, while OCT4 expression remained largely unaffected (Figures 4B and 4C).

To identify downstream effectors responsible for breaking the developmental boundaries of extraembryonic lineages, we analyzed scATAC-seq data from BPSCs, EPSCs, and ESCs. Motif enrichment analysis revealed that LEF1 and TCF3, both being key Wnt pathway effectors, were the most enriched transcription factors in the upregulated accessibility regions of BPSC compared with EPSC or ESC (Figures 4D and 4E). While TCF3 is known to regulate pluripotency genes^40,41^, LEF1 is associated with differentiation and specifically regulates CDX2 expression^42^. We therefore hypothesized that LEF1 drives the conversion of pluripotent stem cells to a bipotent state.

Knockdown of LEF1 under AL conditions significantly reduced the proportion of double-positive cells and downregulated TE lineage gene expression, confirmed by transcriptomics and immunofluorescence showing decreased CDX2 protein levels (Figures 4F, 4G and 4H; Table S2). Similar effects were observed with AS or LY treatments alone (Figures S5C and S5D).

Given the critical role of LEF1 in bipotency, we next examined its impact on the first cell fate decision in embryos. Knockout and knockdown of *Lef1* in fertilized eggs prevented blastocyst formation, with most embryos arresting at the morula stage (Figures S5E and S5F; Table S2). Immunofluorescence revealed significant reductions in CDX2 and SOX2, while transcriptomics indicated impaired expression of TE lineage genes and affected pluripotency genes (Figures S5G-I). These findings suggest that LEF1 plays a crucial role in the first cell fate decision and contributes to lineage stability in early embryonic development.

### Efficient simulation of embryonic development during the early stages of organogenesis using BPSCs

Having established that BPSCs efficiently differentiate into embryonic and extra-embryonic lineages, we next explored their utility in modeling early organogenesis. Notably, BPSCs alone could form blastocyst-like structure (Figures S6A-C). To model post-implantation embryogenesis, we induced BPSCs for 48 hours to generated distinct epiblast- (Epi), trophectoderm- (TE), and primitive endoderm-like (PrE) populations, which were subsequently aggregated using an established in vitro culture (IVC) system^43^ (Figure S6D). Solid spherical embryo-like structures resembling E5.0 embryos emerged after 4 days (Figure S6E). By day 5, Epi-like structure rearranged and formed a lumen-like structure similar to that of E5.5 embryos^39^ (Figures S6F-H).

Next, E5.5-like embryoids were cultured using the *ex utero* culture medium (EUCM) system^44^ (Figure S6D). By day 7, embryoids developed an expanded pro-amniotic-like cavity and initiated primitive streak formation, marked by *Brachyury* (T) expression within the posterior epiblast compartment. This spatial organization recapitulated key features of natural mouse embryos at E6.5–E7.5 stages (Figure S6J). Single-cell transcriptome data showed that the E7.5-like structure was similar to the normal E7.5 embryonic cell lineage (Figure S6K). We cultured the E7.5 gastrulation structures under roller culture conditions for two days and successfully obtained E8.5-like embryoids (Figure S6L). However, the efficiency was relatively low (<1%).

Therefore, we altered the method: for the starting cells (EPSCs or ESCs), one part was transformed into BPSCs and then differentiated into TE-like and Epi-like cells, and another part was directly induced into PrE-like cells (Figures 5A, S7A and S7B). Subsequently, we combined these in microwell plate for embryoid culture under IVC. On the third day, all the aggregates in the well plate were transferred to ultra-low attachment cell culture dishes and cultured with shaking (Figure 5A). By the fourth day, approximately over 75% of E5.5-like structures could be selected for further culture in EUCM (Figures 5A, 5B, S7C and S7D). This efficiency was much higher than that previously reported^24,25^. Starting from the fifth day, the morphologically normal embryoids were subjected to rolling culture until E8.5-like structures emerged on the eighth day, accompanied by regular early cardiac beating (Figures 5B and 5C; Videos S1). Statistical analysis indicated that the number of E8.5-like structures exceeded 16% of all aggregates on day 4 after eight days of cultivation (Figure S7D). The D7 and D8 embryoids developed amnion-like and yolk sac-like structures that correctly encapsulated the embryonic structure (Figures 5B and 5C). Moreover, on the yolk sac-like membrane, regions similar to the blood islands could be observed, along with red pigmentation (Figures 5B and S7C). After dissection, D8 embryoid bodies were observed to possess structures, such as a head fold, beating heart, allantois, and foregut, with a morphology intermediate between E8.5 and E8.75 of normal embryos cultured in EUCM (Figure 5C; Video S2).

**Figure 5.**
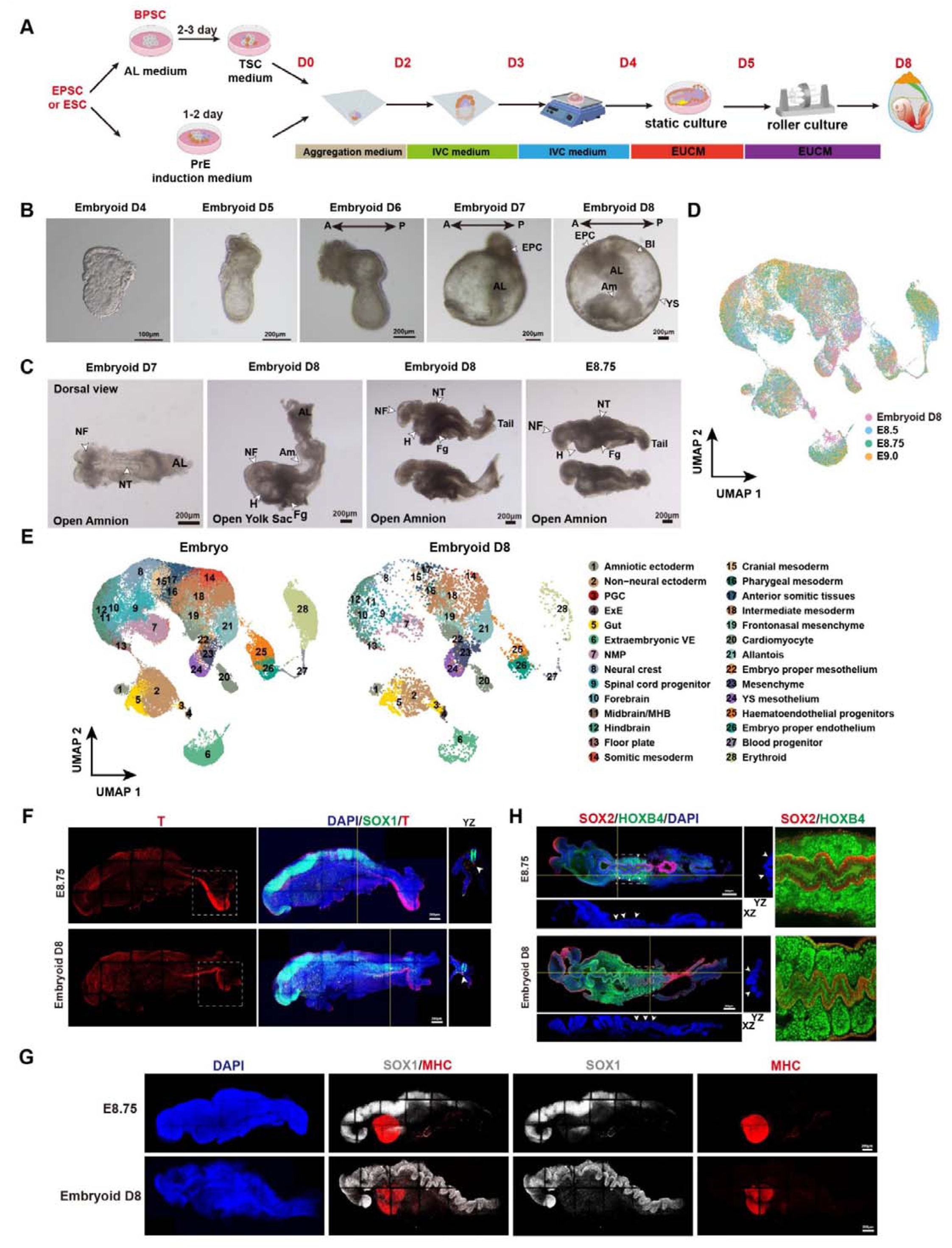
Construction of embryo model by BPSCs. (A) Schematic for constructing embryoids with BPSC. (B) Bright field images of embryoids developing from Day4 to Day8. (C) Bright field images of embryoid D7, embryoid D8, and natural E8.75 embryos. Natural E8.75 embryos were obtained from E7.5 embryos cultured in EUCM for 1.5 days. (D) UMAP of natural E8.5-E9.0 embryos and D8 embryoid derived from BPSCs colored by the stages. (E) UMAP of natural E8.5-E9.0 embryos (left panel) and D8 embryoid derived from BPSCs (right panel). (F) Immunofluorescence staining of natural E8.75 embryo and D8 embryoid. Staining for SOX1 and T. Scale bar, 200 μm. The white dashed box indicates the tail bud and the white arrow indicates the notochord. (G) Immunofluorescence staining of natural E8.75 embryo and D8 embryoid. Staining for SOX1 and MHC. Scale bar, 200 μm. (H) Immunofluorescence staining of natural E8.75 embryo and D8 embryoid. Staining for SOX2 and HOXB4. Scale bar, 200 μm. P, posterior; A, anterior; EPC, ectoplacental cone; Al, allantois; Am, amnion; BI, blood islands; Fg, foregut pocket; H, heart; NF, neural folds; NT, neural tube; YS, yolk sac.

To determine the integrity of the cell lineages in the embryoid bodies, we performed scRNA-seq (Figures 5D and 5E). The results demonstrated that D8 embryoid bodies were highly similar to E8.5-E9.0 embryos, with a relatively complete cell lineage (Figures 5D and 5E). We also statistically analyzed the cell proportions of the main embryonic body, excluding the yolk sac and allantois, and the results showed that the most proportions of lineages were consistent with those of normal embryos (Figure S7E).

Immunofluorescence revealed that, in D8 embryoid, neural folds formed in the head region, and a neural tube developed on the back, gradually extending toward the tail. The expression levels of the neural lineage markers SOX1 and SOX2 followed a pattern of high expression in the head and low expression in the tail. From the images of the three-dimensional reconstructed tissue structure, it was clearly observable that the neural tube near the tail had already closed (Figure 5F). The notochord was distinctly marked by the T protein, which was located at the base of the neural tube (Figure 5F). The enrichment of the T protein in the tail region also indicated the emergence of the tail bud. Through scRNA-seq analysis, the brain could be divided into sub-structures, including the forebrain, midbrain, hindbrain, and floor plate (Figure S7F).

D8 embryoids already exhibited regular heart-beating, and staining for the cardiac marker myosin heavy chain II enabled clear visualization of the early cardiac structure located at the anterior part of the embryoids (Figure 5G). Based on the scRNA-seq data, the heart could be classified into cardiac mesoderm, the first heart field, and the second heart field (Figure S7G). Somites give rise to skeletal muscles, blood vessels, and skin. During the neurula stage, they are arranged in a clump-like structure on both sides of the neural tube. Immunofluorescence staining for HOXB4 showed that D8 embryoid had distinct paired somite structures (Figure 5H).

Surprisingly, using a cell line with the ΔPE-*Oct4*-GFP reporter system, we successfully detected the existence of primordial germ cells (PGCs) in D8 embryoid (Figure S7H). The identical result was observed in scRNA-seq data (Figure 5E). At this stage, PGCs were localized near the allantois and were in the process of migrating towards the anterior part (Figure S7H). scRNA-seq data indicated that the PGCs of D8 embryoids were similar to those in natural embryos, both highly expressing *Dnd1*, *Dppa3*, and *Ifitm3* (Figure S7I). These findings demonstrate that our embryoid model effectively recapitulates key events of natural embryonic development, highlighting its significant potential for modeling early developmental processes.

### Induction of human BPSCs

To further validate the broad applicability of the AL chemical combination in inducing a human BPSC-like state, we assessed its efficacy in promoting bidirectional pluripotent in human naïve embryonic stem cells (nESCs) under 5i culture condition (Figure 6A). Following four days of AL co-treatment, human nESC colonies adopted a more compacted morphology with sharply delineated edges (Figure 6B). Dual nuclear staining followed by flow cytometry analysis revealed a substantial increase in OCT4+/CDX2+ double-positive cells, reaching 48.7% upon AL treatment (Figure 6C). Immunofluorescence further confirmed the nuclear colocalization of OCT4 and CDX2 in these cells (Figure 6D).

**Figure 6.**
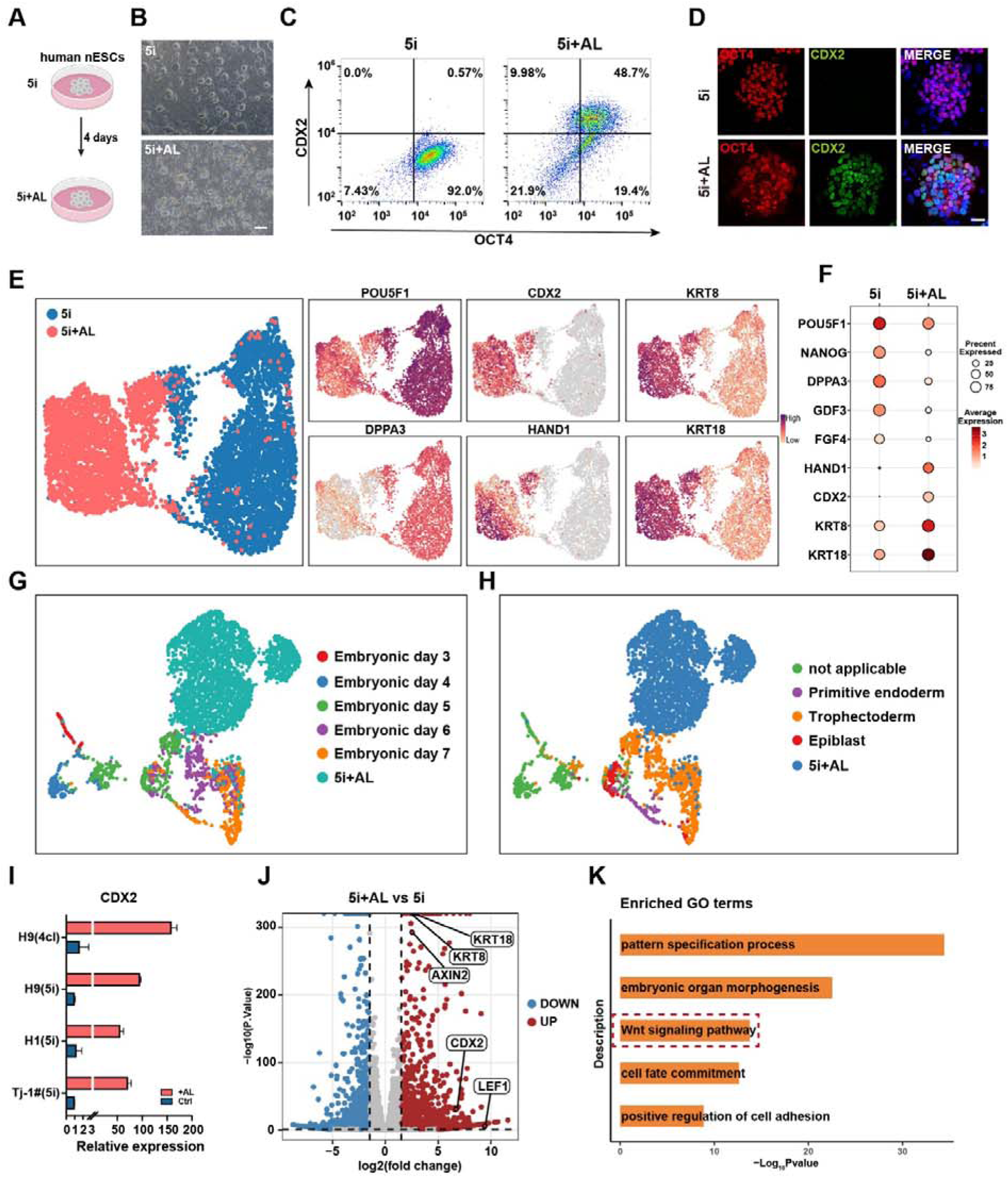
Induction of human BPSCs. (A) Schematic of human naïve embryonic stem cells (nESCs) induction upon AL treatment for 4 days. (B) Morphological changes in human nESCs following AL treatment. Scale bar, 200µm. (C) FACS analysis of human OCT4 and CDX2 expression signals in AL-treated human nESCs. (D) Immunofluorescence staining of OCT4 and CDX2 in AL-treated and 5i human nESCs. Scale bar, 50µm. (E) UMAP visualization of AL-treated and control (Ctrl) human nESCs at day 4. The expression of EPI-specific gene (POU5F1, DPPA3) and TE-specific gene (CDX2, HAND1, KRT8 and KRT18) are displayed on the right side. (F) Bubble plot illustrating EPI-specific gene and TE-specific genes expression in AL-treated and control (Ctrl) human nESCs. (G) Integrated UMAP of single-cell transcriptomes from AL-treated nESCs and natural blastocysts from stage E3 to E7, colored by developmental stage. (H) UMAP embedding of integrated single-cell transcriptomes comprising AL treated nESCs and natural blastocysts. colored by cell lineages. (I) boxplot showing CDX2 expression in AL-treated human nESCs across naïve culture conditions and genetic background. (J) Volcano plot identified differentially expressed genes in AL-treated human nESCs. Red dots denote upregulated genes and blue dots denotes downregulated genes. (K) Gene Ontology (GO) term network analysis of AL-treated human nESCs.

To delineate the molecular dynamics underlying AL-induced bipotency, we performed single-cell RNA sequencing (scRNA-seq) on AL-treated and control human nESCs at day 4 (Figure 6E). Transcriptomic profiling uncovered extensive global reprogramming, characterized by sustained expression of epiblast pluripotency markers (POU5F1, DPPA3) alongside pronounced upregulation of core TE lineage genes (CDX2, HAND1, KRT8, KRT18) (Figures 6E-F). This robust activation of TE-associated transcriptional programs, validated at single-cell resolution, establishes AL as a potent inducer of TE-like states in human nESCs with cross-species applicability.

To further validate the intrinsic identities and lineage commitments of AL-induced human BPSC-like cells, we mapped them onto a single-cell atlas of human embryo development (E3-E7) by integrating with published datasets from natural human pre -implantation embryos. Transcriptomic analysis showed that AL-treated cells aligned most closely with embryonic days 5-6, and integrative analysis confirmed that these cells simultaneously possess both EPI and TE characteristics (Figures 6G, 6H and S8A).

Importantly, this TE priming was consistent and independent of naïve culture conditions or genetic background, as evidenced by the reliable activation of CDX2 across multiple human nESC lines (H9, H1, TJ-1#) (Figure 6I). Furthermore, differential expression (DE) analysis identified genes such as LEF1 and AXIN2 as highly expressed in AL-treated cells, consistent with findings in mouse models (Figure 6J). Gene Ontology (GO) enrichment analysis of AL-upregulated genes revealed significant enrichment in biological processes including pattern specification, embryonic organ morphogenesis, Wnt signaling pathway, and cell fate commitment (Figure 6K).

## Discussion

During early embryonic development, a transient population of CDX2 and OCT4 double-positive cells is observed from the 16-cell to the early blastocyst stage^34^. Previous studies have shown that this unique cell population exhibits both embryonic and extraembryonic developmental potentials^35,37,45^. Capturing and maintaining such cells with bidirectional developmental capabilities *in vitro* could address the issue of low efficiency in differentiating pluripotent and totipotent stem cells into extraembryonic lineages, thereby opening new avenues for fundamental and translational research.

In this study, we successfully captured mouse and human CDX2 and OCT4 double-positive cells via small-molecule screening and designated them as BPSCs. These cells exhibit greater similarity to the 16-cell stage embryo in terms of gene expression and can contribute to both embryonic and extraembryonic tissues *in vivo* and *in vitro*. Unlike the transformation of ESCs and totipotent-like cells into TSCs, which requires the lineage-committing transcription factor CDX2 or long-tine culture, BPSCs can attain a high percentage of TSCs with short-term culture in TSC culture medium. Compared to previously reported extended pluripotent stem cells and totipotent-like stem cells, BPSCs demonstrated efficient differentiation into all three early embryo lineages. *In vitro* differentiation experiments confirmed that *Oct4*+/*Cdx2*+ cells are the main source of TE-like cells, and PrE cells are mainly differentiated from OCT4 single-positive cells. This finding suggests that BPSCs represent a more functionally and ecologically advanced cell type than pluripotent ESC and TSC lines.

The successful capture of BPSCs relies on the small molecules LY and AS, which function as inhibitors of GSK3 and FOXO1, respectively. The WNT pathway has been reported to activate CDX2 expression in mouse and rat embryonic stem cells^42,46^. We increased the concentration of the classic WNT pathway activator CHIR99021 to 10 μM and obtained the same effect as that of LY (10 nM). Previous studies demonstrated that FOXO1 plays a crucial role as a transcription factor in maintaining the stemness and self-renewal capabilities of ESCs. Additionally, deletion of *Foxo1* has been shown to affect the expression of multiple pluripotency genes, such as *Oct4* and *Sox2*^47,48^. Increasing concentrations of AS initially facilitated the maintenance of OCT4, and further increases in concentration led to the emergence of cells that were positive for OCT4 and CDX2. The above results contrast with the trans-differentiation process observed in previous studies, which involved *Oct4* downregulation and *Cdx2* upregulation^42^. In this study, we were surprised to find that AS primarily activates CDX2 expression through the WNT signaling pathway, rather than FOXO1. This suggests a potential close relationship between these two pathways, warranting further investigation in future studies.

The emergence of embryo-like structures from BPSCs circumvents the scarcity of embryos for developmental studies and black-box characteristics of the implantation process, thereby promoting the field of developmental biology. Previously, the construction of embryo-like structures required complex cell combinations and suffered from low efficiency. The present study demonstrates that BPSCs can achieve E5.5–E8.5-like structures by changing the culturing system.

In conclusion, we successfully captured a unique bidirectional developmental cell state (BPSCs), distinct from totipotent and pluripotent states, through the combination of the small molecules AS and LY. This study provides a promising approach for lineage development and the construction of embryo-like structures.

## Supporting information

Supplementary Video 1. Embryoids D8 in yolk sac with beating heart

Supplementary Video 2. Embryoids D8 with beating heart after amniotic membrane cut

Supplementary Table 1. Small molecules and targets

Supplementary Table 2. sgRNA & siRNA sequences

## Acknowledgments

This work was primarily supported by the National Key R&D Program of China (2021YFA1102900 and 2022YFC2702200) and the National Natural Science Foundation of China (32488101, 32330030, 82301921, 32070823, 92168205, 32270909, 82201881, 92168205, 31871448, 31820103009, 82022027, 32300684 and 32400677). This work was also supported by the key project of the Science and Technology of Shanghai Municipality (19JC1415300 and 21JC1405500), the Natural Science Foundation of Shanghai (21ZR1450700), China Postdoctoral Science Foundation (2023M732660, 2023M732661), the Postdoctoral Fellowship Program of CPSF (GZB20230523), Shanghai Pilot Program for Basic Research, Shanghai Post-doctoral Excellence Program (2022521, 2022551), Shenzhen Medical Research Fund (B2402013), the Fundamental Research Funds for the Central Universities (Grant Nos. 22120240435), Peak Disciplines (Type IV) of Institutions of Higher Learning in Shanghai and Ministry of Education for their support.

## Author contributions

S.G., W.L., Y.W. conceived the project and provided mentoring; K.L., D.B and Y.B. designed and performed the most experiments; Z.Y. provided bioinformatics support; R.J and K.W performed compound library screening; K.L., D.B., Z.Y., wrote original draft; K.L., D.B., Z.Y., W.L., J.Y., S.G., performed writing – review & editing; J.X.,Y.S., B.D., X.M., Z.N., S.Y., Y.L., X.L., Y.J., Y.Z., Y.Z, Y.L., C.X., S.L., S.L., J.C., J.Y., X.K., Y.Z., H.W. assisted with the experiments.

## Supplemental information titles and legends

Supplementary Table 1. Small molecules and targets

Supplementary Table 2. sgRNA & siRNA sequences

Supplementary Video 1. Embryoids D8 in yolk sac with beating heart

Supplementary Video 2. Embryoids D8 with beating heart after amniotic membrane cut

## Methods

### RESOURCE AVAILABILITY

#### Lead contact

Further information and requests for resources and reagents should be directed to and will be fulfilled by the Lead Contact, Shaorong Gao.

#### Materials availability

This study did not generate new unique reagents.

#### Data and code availability

The bulk RNA-seq, single cell RNA-seq and single cell ATAC-seq data generated during this study are available at Genome Sequence Archive of the National Genomics Data Center (https://ngdc.cncb.ac.cn/gsa/): PRJCA027661.

### EXPERIMENTAL MODEL AND STUDY PARTICIPANT DETAILS

#### Animals

All animal experiments were performed following the Guide of experimental animal use guide and approved by Tongji University. All animal were housed and bred in a specific pathogen-free (SPF) animal research facility at Tongji University, Shanghai, China. The C57BL/6, ICR, 129 mice at 6-8 weeks old were purchased from Beijing Charles River (China). B6.Cg-Tg(CAG-GFP)Smoc mice with stable fluorescent gene expression in all organs were obtained from Shanghai Model Organisms Center (China). TgOG2 mice with △PE-*Oct4*-GFP (This article is referred to as *Oct4*-GFP) fluorescent reporter were generous gift from Dr. Jeff R. Mann, University of Melbourne, Victoria, Australia^56^. CDX2-mCherry fluorescent reporter mice were constructed by Professor Jiayu Chen, Tongji University, Shanghai, China^55^. The *Oct4*-GFP and CDX2-mCherry double reporter mice were obtained by crossing TgOG2 mice (female, 6-8 weeks old) with CDX2-mCherry fluorescent reporter mice (male, 8-10 weeks old). F1 hybrids were generated between B6.Cg-Tg(CAG-GFP)Smoc (male, 8-10 weeks old) and ICR (female, 6-8 weeks old) and served as a source of cell lines for chimera. Chimeric donor embryos were obtained from C57BL/6 females aged 6–8 weeks. Pseudopregnant recipients were ICR females aged 8-10 weeks.

#### Cell lines

Mouse ESC/EPSC with *Oct4*-GFP and CDX2-mCherry double-fluorescent reporter were derived from E3.5 blastocyst (*Oct4*-GFP and CDX2-mCherry double-fluorescent reporter mice); WT mouse ESC/EPSC were derived from E3.5 blastocyst (129 female mice crossed with male OG2 mice); ESC/EPSC with persistent CAG-GFP fluorescence were derived from E3.5 blastocyst (ICR female mice crossed with male B6.Cg-Tg(CAG-GFP)Smoc).

### Methods details

#### Culture of Mouse Stem Cells

Mouse fibroblasts were treated with mitomycin (MMC; Sigma–Aldrich, M0503) as feeder for culturing mouse stem cells, and only a few experiments requiring feeder free. All mouse stem cells were cultured at 37°C in a humidified incubator with 5% CO_2_. Mouse stem cells were replaced culture medium every day or every other day, and were passaged using 0.05% trypsin-EDTA when cells at higher densities (80% confluence).

ESC medium (ESM) was a basal medium used to culture ESC which containing 1 mM L-glutamine (Millipore, 25030–081), 0.1 mM β-mercaptoethanol (GIBCO, 21985–023), 1× nonessential amino acid (Millipore, TSM-001-C), 1× penicillin–streptomycin (Thermo Fisher, 15140122), and 1× nucleosides (Millipore, ES-008-D) and 15% (v/v) fetal bovine serum (FBS) (HyClone, SH30070.03) in DMEM (Sigma–Aldrich, D5671). In this research, ESC was cultured in ESM supplemented with 2i/LIF (1 mM MEK inhibitor PD0325901 Selleck 1036S; 3 mM GSK-3 inhibitor CHIR99021 Selleck S2924 and 10 ng ml^-1^ rmLIF) or with LIF (10 ng ml^-1^ recombinant mouse LIF; Millipore, ESG1107).

EPSC were cultured in EPSC medium which has been described in published study^68^. Preparation of N2B27 basal medium: DMEM/F-12 (GIBCO, 11330–032) and neurobasal medium (GIBCO, 21103–049) were mixed in a 1:1 ratio, then added with 1× nonessential amino acid, 1 mM L-glutamine, 0.1 mM β-mercaptoethanol, 5% KnockOut Serum Replacement (GIBCO 10828–028), 0.5× N2 supplement (Invitrogen, 17502–048) and 0.5× B27 supplement (Invitrogen 17504–044). EPSC medium preparation was completed by adding 3 mM CHIR99021, 2 mM (S)-(+)-dimethylene maleate (Tocris, 1425), 2 mM minocycline hydrochloride (Selleck, s4226) and 10 ng/mL recombinant human LIF (Millipore, LIF1050) to N2B27 basal medium.

BPSC were cultured in N2B27 basal medium supplemented with 10 nM LY2090314 (MedChem Express HY-16294) and 0.6 μM AS1842856 (MedChem Express HY-100596).

TSC were cultured in TSC medium (TSM) as described in published study^7^. Preparation of TSM basal medium: RPMI 1640 medium added with 20% (v/v) FBS, 1× GlutaMAX (Thermo Fisher, 35050–061), 0.1 mM β-mercaptoethanol, 1× sodium pyruvate (Sigma–Aldrich, S8636) and 1× penicillin-streptomycin. TSM was based on TSM basal medium supplemented with 25 ng/mL FGF4 (Sino Biological 16043-HNAE-100) and 1μg/mL Heparin (Sigma-Aldrich H3149-500KU).

PrE induction medium was composed of N2B27 basal medium supplemented with 2.5 μM 1-Azakenpaullone (Selleck Chemicals, S7193), 100 ng/mL recombinant human FGF4 (PeproTech, 100–31), 0.2 μM TTNPB (Selleck Chemicals, S4627), and 1 mM 8Br-cAMP (Selleck, S7857).

#### Chemical induction of BPSC

##### Chemical induction of BPSC from mouse ESC/EPS

Mouse ESC/EPSC with *Oct4*-GFP and CDX2-mCherry double-fluorescent reporter were cultured with N2B27 basal medium supplemented with 10 nM LY2090314 (MedChem Express HY-16294) and 0.6 μM AS1842856 (MedChem Express HY-100596) for 2-3 days. BPSC with *Oct4*-GFP and CDX2-mCherry double positive were detected with flow cytometry (FACS) and fluorescence microscope. ESC/EPSC with persistent CAG-GFP fluorescence were cultured with BPSC medium for 3 days and as a source for chimera.

##### Derived BPSC from morula

BPSC were derived from morula (16- to 32-cell) which obtained from *Oct4*-GFP and CDX2-mCherry double-fluorescent reporter mice. Cultured with BPSC medium, the karyotype can be identified after several passages.

#### Culture of Human Stem Cells

Naive human PSCs were maintained in 5iLAF medium, which consists of naïve basal medium containing DMEM/F12:Neurobasal (1:1) (Thermo Fisher), 1% N2 supplement (Thermo Fisher), 2% B27 supplement (Thermo Fisher), 0.5% KnockOut SR (Thermo Fisher), 1% nonessential amino acids (Millipore), 2 mM GlutaMAX (Millipore), and penicillin-streptomycin (Millipore) with 1 μM PD0325901 (Selleck), 0.5 μM SB590885 (Selleck), 1 μM WH-4-023 (Selleck), 1 μM IM12 (Selleck), 10 μM Y-27632 (Selleck), 20 ng/ml activin A (PeproTech), 20 ng/ml human LIF (Millipore), 8 ng/ml bFGF (PeproTech), and 50 μg/ml BSA (Sigma-Aldrich).

4CL medium composed of 1:1 mix of Neurobasal medium (Gibco, 21103049) and Advanced DMEM/ F12 (Gibco, 12634028) supplemented with 1% N2 (Gibco, 17502048) and 2% B27 (Gibco, 17504044), 1mM sodium pyruvate (Corning), non-essential amino acids (Corning), 1mM GlutaMAX (Gibco), penicillin–streptomycin (Millipore), 10 nM DZNep (Selleck), 5 nM TSA (Selleck), 1 μM PD0325901 (Selleck), 5 μM IWR-1 (Selleck), 20 ng/ml human LIF (Peprotech), 20 ng/ml activin A (Peprotech, 120-14E), 50 μg/ml L-ascorbic acid (Sigma) and 0.2% (v/v) Matrigel.

Naïve cells were passaged with Accutase (Sigma-Aldrich) every 4-7 days. All human cell lines were cultured in 5% O_2_ and 5% CO_2_ at 37 [. Mycoplasma were routinely tested every week. Human ESC lines were used in accordance with the ethical approvals obtained from the Biological Research Ethics Committee of Tongji University.

10 nM LY (MedChem Express) and 0.6 μM AS was supplemented in 5iLAF or 4CL for induction.

#### Differentiation of BPSC

For the autonomous differentiation of BPSC

BPSCs were obtained by culturing BPSC in plate for 2 days with inducing-factor-free TSM basal medium.

For the induced differentiation of BPSC

BPSC differentiation with TSM with 25μg/mL FGF4 and 1μg/mL Heparin for 3 days and collected CDX2-mCherry positive or CDCP1 (R&D Systems AF4515-SP) positive cells with FACS. Multi-directional dividing cells were induced with TSM for 2 days and used to chimera.

#### Chimeras and microinjection of differentiated cells (BPSC differentiation with TSM) and ESC into embryos

Chimeras were constructed by microinjection of differentiated cells and ESCs into normal donor embryos at 8-cell stage. The constructed chimeras were cultured in G-1 PLUS (Vitrolife 10128) medium and transferred into uterus of 2.5 days post pseudopregnant recipients at E3.5 blastocyst stage.

#### Tetraploid complementation experiment

Recipient embryos at the 2-cell stage were collected from the oviducts of mated ICR females and cultured in G-1 PLUS medium under mineral oil conditions at 37°C and 5% CO_2_ using the microdrop method. Electrofusion produced tetraploid embryos at the late 2-cell stage. The tetraploid embryos were cultured in G-1 PLUS medium until aggregation. BPSC colonies were treated with 0.05% trypsin-EDTA for about 4 minutes in a 37°C cell culture incubator to obtain single-cell suspensions.

10-15 single cells were transferred to a depression well of an aggregation plate. The zona pellucida of tetraploid embryos at the “4-cell stage” was digested with 20 mg/mL PE (Pronase E; Sigma, #P8811), and two embryos were placed as a group in a depression well. Aggregated embryos were cultured in G-1 PLUS medium until the blastocyst stage and then transferred into the uterus of a pseudopregnant recipient at 2.5 dpc.

#### Cell transfection

For small interfering RNA (siRNA) transfections, Lipofectamine RNAiMAX reagent (Invitrogen, Catalog No. 13778100) was first diluted in Opti-MEM reduced-serum medium (GIBCO, Catalog No. 31985070). The diluted reagent was then mixed with siRNAs that had been separately diluted in the same medium. This mixture was incubated at room temperature for 15 minutes to allow the formation of transfection complexes.Following the incubation, the transfection complexes were added dropwise to the cell cultures. Cells were maintained under this culture conditions for 48 hours to allow sufficient time for siRNA-mediated gene knockdown. After the 48-hour incubation period, cells were harvested for downstream assays. The sequences of the siRNAs used in these experiments are listed in Table S1.

#### Knockdown (KD) and Knockout(KO) in embryos

Three siRNAs and three sgRNAs targeting *Lef1* were pooled and diluted to a final concentration of 20 μM (siRNA) and 75 ng/μl (sgRNA) for microinjection. Zygotes derived from B6D2F1-crossed female mice were microinjected with the mixed siRNA/sgRNA solution using a Piezo-driven micromanipulator. Following microinjection, embryos were cultured in G-1 PLUS medium (Vitrolife) under controlled conditions (5% CO_2_ at 37°C).

#### Construction of embryo-like structures with IVC and EUCM culture system

##### Construction of embryo-like cells by sequential induction of TS and PrE

We constructed embryo-like structures with IVC and EUCM culture system followed previously reported^24,25,43^. BPSC was induced in TSM for one day and then induced to PrE for another day. The cells were then digested into single cells and planted in aggrewell at a rate of 31,200 cells per well. On day one, use FM plus 10 μM Y-27632. Wash with fresh FM twice the next day, and continue to culture with fresh FM. On the third day, 1.5 ml of IVC2 was replaced per well, and on the fourth day, 2ml of IVC2 was replaced. On the fifth day, the embryoids were transferred into ultra-low adhesion 6-well and continued to be cultured with IVC2 for one day, and then the embryoids with good shape were transferred into EUCM for two days. For the next 2 days, the embryoids were cultured in roller culture conditions (rotating at 30 revolutions per minute) with fresh EUCM replaced daily and extra 3 mg/L glucose added on the last day.

##### Construction of embryo-like structures by separate induction of TS and PrE

First, the starting cells EPSC or ESC were divided into two parts. One part was first converted into BPSCs and then transferred to TSM for culture for two days. The other part was cultured in PrE induction medium for one (EPSC) or two (ESC) days. The cells were then digested into single cells and planted in aggrewell at a rate of 43,200 cells (35200 cells of TS induction plus 8000 cells of PrE induction) per well. On the first day, use a mixed medium (1:1) of FM and N2B27 basal medium plus 10 μM Y27. The next day, continue to culture with fresh mixed medium without Y27. On the third day, 1.5 ml of IVC2 was replaced per well. On the fourth day, all the aggregates were transferred into ultra-low adhesion 6-well under shaking condition (rotation 70 rpm/min) and continued to be cultured with IVC2 (3 ml per well) for one day. On the fifth day, the embryoids with good shape were transferred into EUCM under static culture conditions for one days. For the next 3 days, the embryoids were cultured in roller culture conditions (rotating at 30 revolutions per minute, Eastmo biotech Ltd, WEC001-GT4) with fresh EUCM replaced daily and extra 3 mg/L glucose added on the last day.

FM was composed of DMEM containing 15% (v/v) fetal bovine serum (FBS), 1 mM L-glutamine, 1× nonessential amino acid, 1× penicillin–streptomycin and 1× sodium pyruvate. IVC2 was composed of CMRL 1066 (Thermo Fisher Scientific 11530037) containing 20% (v/v) FBS, 1 mM sodium pyruvate, 1× penicillin-streptomycin, 2 mM L-glutamine, 1× N2 supplement and 0.25× B27 supplement. EUCM was composed of 25%(v/v) DMEM (Sigma–Aldrich, D5671) or DMEM/F12 (GIBCO, 11330–032), 25%(v/v) human adult blood (Sigma-Aldrich, H4522-100ML), 50% rat serum (Charles River), 1 mM sodium pyruvate, 1× nonessential amino acid, 1× penicillin-streptomycin and 2 mM L-glutamine.

#### Generation of BPSC-blastoids

The 6000-12000 cells were placed in one well of a 24-well Aggrewell-400 (STEMCELL Technologies, 34415) plate cultured with blastoid medium. Blastoid medium consisted of 25% TSC basal medium, 25% (v/v) N2B27 basal medium, and 50% (v/v) KSOM (Aibei Biotechnology, M1430) or G-1 PlUS, supplemented with 2 µM ROCK inhibitor Y-27632 (Selleck, S049), 12.5 ng/mL recombinant human FGF4, 0.5 µg/mL heparin (Sigma–Aldrich, H3149), 3 µM CHIR99021, 5 ng/mL recombinant human BMP4 (PeproTech, 12–05ET), and 0.5 µM A83–01 (Axon Medchem, 1421). The medium was replaced with fresh culture medium without Y-27632 the next day.

#### Immunofluorescence

Before Immunofluorescence, all cell clones and embryos need to fixed with 4% paraformaldehyde (Servicebio, China) at 4°C for 24h. After fixation, embryos including morula, blastocysts and E6.5, cell colonies, E5.5-like embryoid were incubated with permeabilisation reagents for 30 minutes at room temperature. Permeabilisation reagents were compounded with 0.3% Triton X-100 (Sigma–Aldrich, 93443) in DPBS (Gibco). Then, cell colonies were blocked with blocking reagent containing 3% bovine serum albumin (BSA, MP Biomedicals) in DPBS for 1 hours at room temperature. Unlike cell colonies, embryos and E5.5-like embryoid need to be permeabilized and blocked with 0.5% Triton X-100 and 3% BSA in DPBS for 2 hours at room temperature. Primary antibodies were diluted in blocking reagent and incubated with samples overnight at 4°C. After 3 times washed with DPBS containing 0.01% Triton X-100, secondary antibodies were diluted in 3% BSA-DPBS and incubated with samples at room temperature for 2 hours. Nuclei were stained with 4’,6-Diamidino-2-phenylindole (DAPI) (Invitrogen D3571) for 15-20 minutes at room temperature. After nuclei stained, samples were washed samples were washed using DPBS. Samples completed immunofluorescence were photographed and processed with ZEISS LSM 880 confocal microscope.

##### For whole-mount immunostaining of embryoids D8

We cut the embryoids out of the amnion, placed them in 4% paraformaldehyde, and fixed them overnight at 4°C. Before incubating the primary antibody, we used the Animal Tissue Optical Clearing Kit (Beyotime, P0112L) to clear the embryoids. Then, we performed immunofluorescence staining according to the above method, and doubled the incubation time of the secondary antibody and DAPI. Samples completed immunofluorescence were photographed and processed with Olympus FVMPE-RS microscope.

The primary antibodies used and dilutions were as follows: mouse anti-CDX2 (1:400; Bio-Genex, MU392A-UC), anti-OCT4 (1:200; Santa Cruz, sc-5279), anti-GATA6 (1:200; R&D Systems, AF1700), anti-SOX2(1:200; R&D Systems, AF2018-SP), anti-Brachyury(1:200; R&D Systems, AF2085-SP), anti-TFAP2C (1:200; Santa Cruz, sc-12762), anti-SOX17 (1:200; R&D Systems, AF1924-SP), anti-SOX1 (1:200; Cell Signaling Technology,4194S), anti-MHCII (1:200; R&D Systems, MAB4470-SP), anti-HOXB4 (1:100; Abcam,ab133521), anti-TPBPA (1:500; Abcam, ab104401), Secondary antibodies were purchased from Thermo Fisher’s Alexa Fluor® and diluted at 1:200.

#### Fluorescence positive cell sorting

Cells were digested by 0.05% trypsin-EDTA and washed with DPBS supplemented with 2% FBS. After stained with anti-CDCP1 (1:50; R&D Systems, AF4515-SP) or anti-PDGFRα (1:200; Abcam, ab203491) antibodies, cells were washed and resuspended with DPBS supplemented with 2% FBS.

Cell with *Oct4*-GFP and CDX2-mCherry double-fluorescent reporter were unstained and subjected to FACS analysis. All cells were analyzed and collected with MoFlo XDP cell sorter (Beckman Coulter).

#### Compound library screening

A high-throughput screening was conducted using *Oct4*-GFP and CDX2-mCherry double-labeled, stably expressing extended pluripotent stem cells (EPS). These EPSCs were seeded into 384-well plates 24 hours after feeder cells plating. After allowing 12 hours for cell attachment, a total of 1859 compounds from the Selleck Chemicals library (USA) were added to the cells at a final concentration of 1 µM. Each compound was tested in triplicate wells.

Following a 72-hour incubation period, cells were fixed with 4% paraformaldehyde (PFA) for 20min. The nuclei were subsequently stained with DAPI. High-content imaging was performed using the Operetta CLS system (PerkinElmer, USA), and image analysis was conducted using Harmony 4.0 software (PerkinElmer, USA).

#### Bulk mRNA-seq library

Cells in 35 mm dish at 80% confluence were digested with 0.05% trypsin-EDTA. After washed with DPBS, cells were collected in a 1.5 mL tube containing 1 mL TRIzol (Takara Bio, 9109). After vortexed 5 min, cells can be transferred to −80 °C until extract RNA. Phenol chloroform extraction method was used to extract total RNA. KAPA Stranded mRNA-Seq Kit (KAPA, KK8421) were used to construct bulk mRNA-seq libraries. Constructed mRNA-seq libraries were sequenced on Illumina Novaseq 6000 with paired-end of 150 bp. Nanjing Jiangbei New Area Biomedical Public Service Platform performed sequencing and quality control of the constructed libraries.

#### Preparation of single cells for Single cell RNA sequencing (scRNA-seq)

Cells were digested with 0.05% trypsin-EDTA and wash 3 times with DPBS supplemented 0.04% BSA. Embryoids were digested with a mixture of accumax (Sigma-Aldrich A7089-100ML) and Collagenase (Thermo Fisher Scientific 17104019) in a 1:1 ratio for 10-20 min at room temperature. Samples for scRNA-seq should meet a cell viability of more than 80% and clustering rate of less than 20%. Single cell RNA libraries were constructed by 10× Genomics Chromium next GEM Single cell 3’ kit v3.1(16 rxns PN-1000268) and MGI DNBelab C RNA library V2.0. Libraries were sequenced on Illumina Novaseq Xplus and MGI DNBSEQ-T7.

#### Bulk RNA-seq analysis

For the bulk RNA-seq data analysis, we utilized the software Trim Galore v0.6.10 to trim adaptors. Subsequently, STAR 2.7.10b was employed to align the data against the mouse reference genome mm10. The gene expression matrix was generated using FeatureCounts v2.0.3. Differentially expressed genes were calculated and Principal Component Analysis (PCA) was performed using the DESeq2 v3.4.1 package. Genes were considered significant if |log2(fold change)| > 0.5 and adjusted P < 0.05. For the GO analysis, we utilized Metascape (https://metascape.org/gp/index.html) following the provided instructions^69^, or using clusterProfiler R package with default parameters^70^. Significant GO terms were visualized with the ggplot function within the ggplot2 package in R.

#### scRNA-seq analysis

To obtain the gene expression matrix, scRNA-seq FASTQ files were analyzed using either CellRanger v7.1.0 (https://support.10xgenomics.com/single-cell-gene-expression/software/overview/wel come) or DNBC4tools v2.1.1 (https://dnbc4tools.readthedocs.io/zh/latest/index.html). Cells with low gene detection (nFeature), extreme total RNA counts (nCount), or high mitochondrial gene content (pctMT) were excluded. For dimensional reduction analysis, the scRNA-seq data were normalized and integrated using the Seurat package v4.3.0 following standard pipelines^65^. The “RunHarmony” function from the Harmony package v0.1.1 was applied to remove batch effects^66^. Dimensional reduction was achieved through Uniform Manifold Approximation and Projection (UMAP).

#### scATAC-seq analysis

The scATAC-seq and FASTQ files were analyzed using Cellranger ATAC v2.1.0 (https://support.10xgenomics.com/single-cell-atac/software/overview/welcome). The package Seurat v4.3.0 and Signac v1.11.0 were used for quality control, data integration and dimensional reduction and differentially accessible peak analysis according to the standard pipeline^63^. The peak calling was performed using the function “ CallPeaks” in Signac package based on MACS2 v 2.2.7.1. Motif enrichment analysis was performed using the HOMER software package v4.11.1 with the findMotifsGenome.pl script^64^. For the transcription factor footprinting analysis, the motif information was obtained from the package JASPAR2020, and footprinting information was gathered by the function “Footprint” in Signac following the official documentation.

#### ChIP-seq analysis

The ChIP-seq analysis was performed following the standard pipeline^71,72^. Adaptor trimming was conducted using Trim Galore v0.6.10. Next, the software bwa v0.7.17 was employed to map the reads to the mouse reference genome mm10^73^. The tool “bamCompare” from deepTools v3.5.4 was utilized to convert BAM files into BigWig files^61^. To eliminate erroneous regions with high ChIP-seq signal, we downloaded the blacklist region BED file from https://github.com/Boyle-Lab/Blacklist/ and filtered these regions before converting BAM files into BigWig files^74^. The peak calling was performed by the software MACS2 v 2.2.7.1. Additionally, IGV v2.16.0 was used to view the ChIP-seq signal of specific genes^62^.

#### Public data collection

To establish the developmental trajectory of early mouse embryos, we collected bulk RNA-seq datasets GSE66582^50^, CRA003985^49^, and scRNA-seq datasets GSE45719^52^, GSE123046^39^ from the Gene Expression Omnibus or Genome Sequence Archive of the National Genomics Data Center. The scRNA-seq of mouse E8.5-E9.0 embryos and human E3-E7 embryos were downloaded from the dataset E-MTAB-11763^53^ and E-MTAB-3929^54^ on ArrayExpress, respectively. To compare our BPS cell line with other cell lines, we also downloaded several cell line datasets, including GSE201751^19^ (ESC/EPSC, scRNA-seq/scATAC-seq) and CRA004846^51^ (ESC/EPSC, bulk RNA-seq/chip-seq). All of these data were downloaded and analyzed following the same procedures as our data.

#### Data and code availability

The bulk RNA-seq, chip-seq, single cell RNA-seq and single cell ATAC-seq data generated during this study are available at Genome Sequence Archive of the National Genomics Data Center (https://ngdc.cncb.ac.cn/gsa/): PRJCA027661.

---

**Figure S1.**
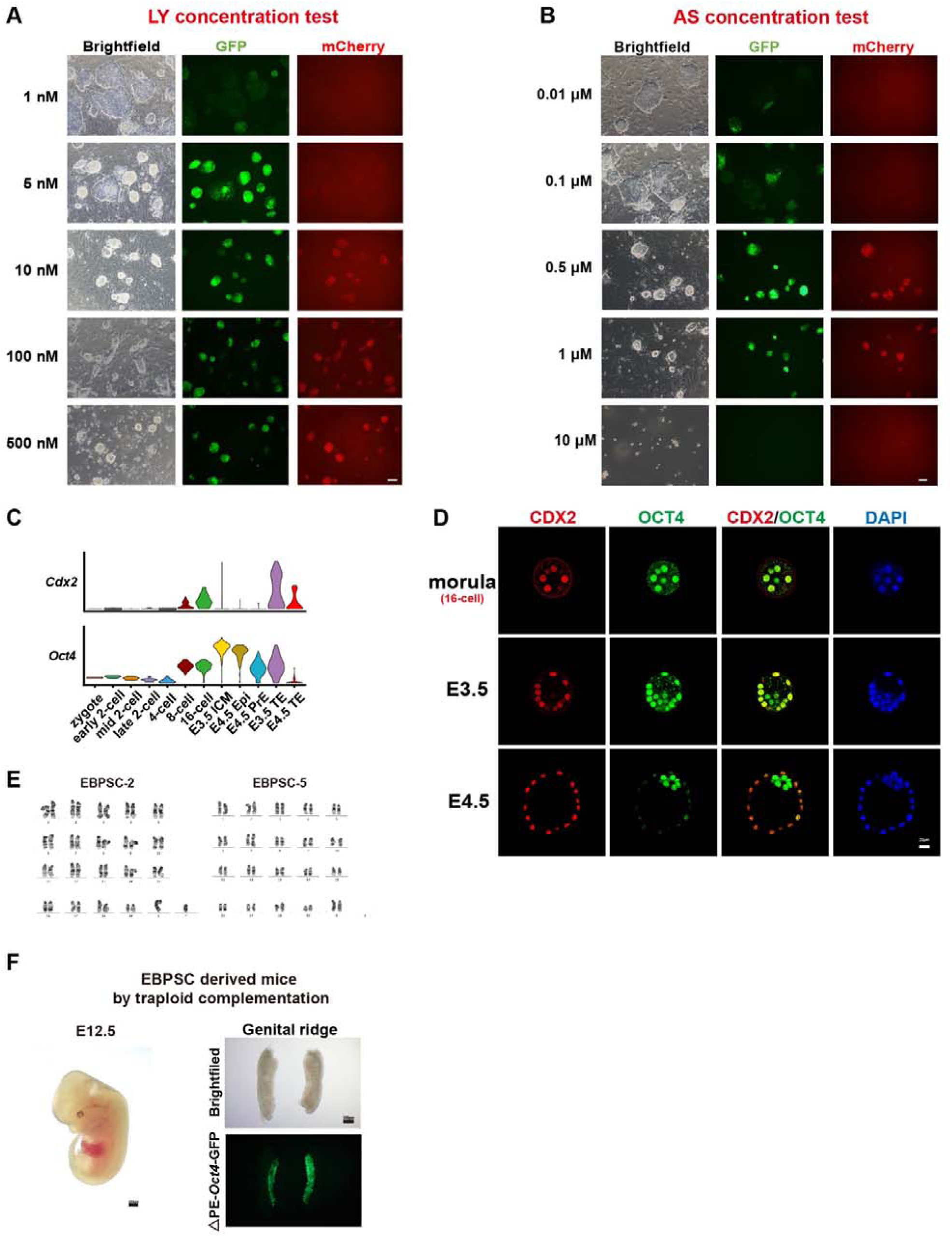
Chemical induction of bidirectional pluripotent stem cells (BPSCs) with double-positive of OCT4 and CDX2. Related to Figure 1. (A) Concentration test of cells treated with LY. Representative morphological images and *Oct4*-GFP, CDX2-mCherry fluorescence were showed. Scale bar, 100 μm. (B) Concentration test of cells treated with AS. Representative morphological images and *Oct4*-GFP, CDX2-mCherry fluorescence were showed. Scale bar, 100 μm. (C) Violin plot showing representative gene expression (*Cdx2*, *Oct4*) in preimplantation embryos from zygotes to E4.5 blastocysts. (D) Immunofluorescence staining of the morula (16-cell stage, top), E3.5 blastocyst (middle) and E4.5 blastocyst (bottom). Staining for CDX2 and OCT4. Scale bar, 20 μm. (E) The karyotype analysis of EBPSC (derived from morula in AL medium). (F) Brightfield image of tetraploid complemented embryo and genital ridges at day E12.5. △PE-*Oct4*-GFP fluorescence reveals the location of PGCs.

**Figure S2.**
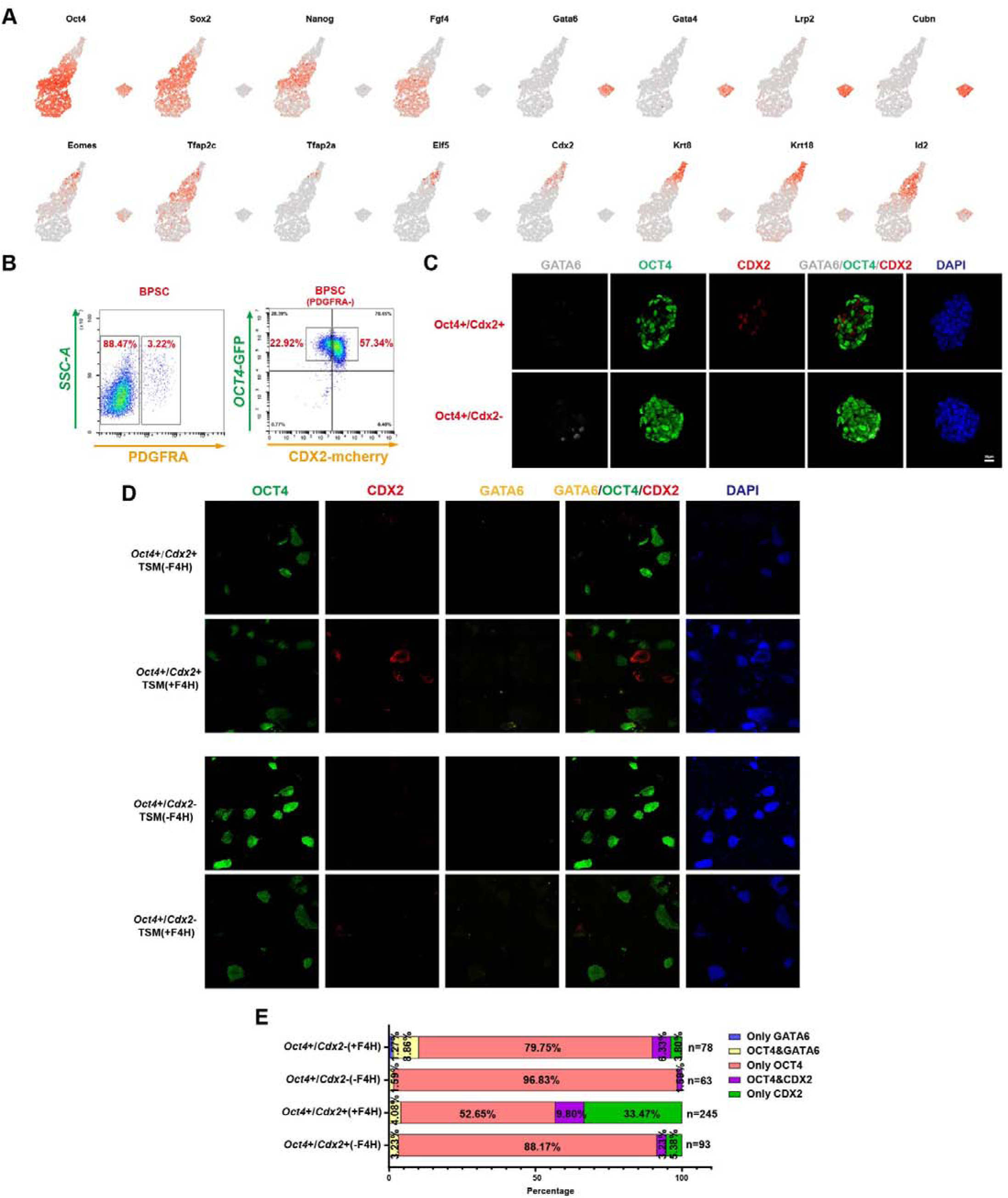
BPSCs exhibit efficient spontaneous differentiation into TE-like cells. Related to Figure 2. (A) Representative gene expression of Epi lineage, TE lineage, and PrE lineage in E4.5 embryo cells, BPSC (derived from EPSC) after 1 or 2 days of spontaneous differentiation, and EPSC after 2 days of spontaneous differentiation. (B) Flow cytometry was used to sort single positive and double positive cells and exclude PDFFRA positive cells. (C) Immunofluorescence staining of Oct4+/Cdx2+ and Oct4+/Cdx2-cells after 2 days of spontaneous differentiation. Staining for OCT4, CDX2 and GATA6. Scale bar, 20 μm. (D) Immunofluorescence staining of differentiation cells with or without FGF4 & Heparin. Staining for OCT4, CDX2 and GATA6. Scale bar, 20 μm. (E) Stacked bar plot showing the percentages of Oct4+/Cdx2+ and Oct4+/Cdx2-cell differentiation with or without FGF4 & Heparin.

**Figure S3.**
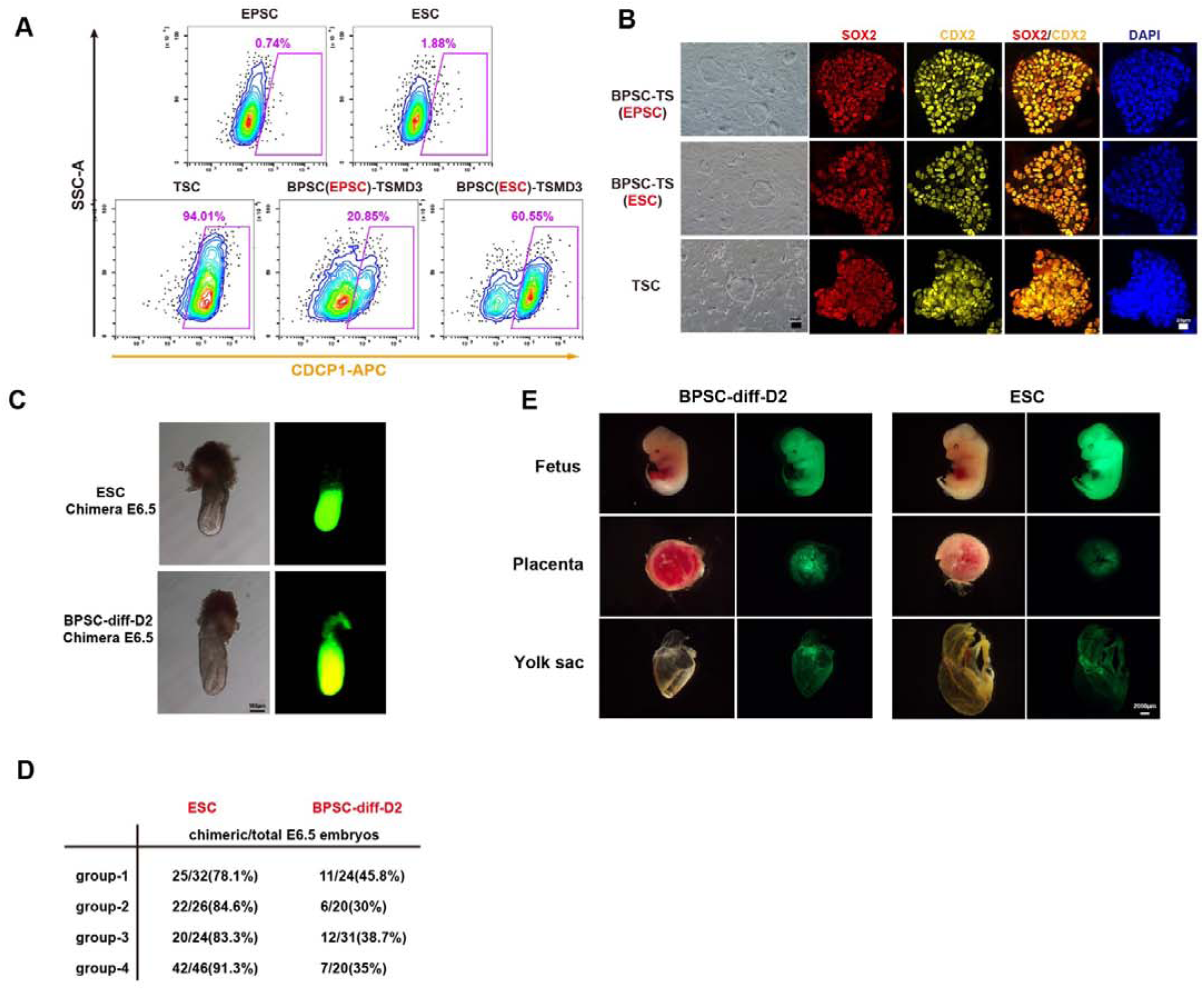
BPSCs exhibit efficient spontaneous differentiation into TE-like cells. Related to Figure 2. (A) Representative FACS analysis of the percentages of CDCP1 positive cells in EPSC, ESC, TSC, BPSC(EPSC)-TSMD3 (EPSC-derived BPSC induced with TSM for 3 days) and BPSC(ESC)-TSMD3 (ESC-derived BPSCs induced with TSM for 3 days) (B) Immunofluorescence staining of BPSC-TSC (EPSC), BPSC-TSC (ESC) and TSC (derived from embryo). Staining for SOX2 and CDX2. Scale bar, 20 μm. (C) Representative morphological images of E6.5 chimeric embryo produced using the above method (Figure 2F). (D) E6.5 Chimera embryo formation ratio of differentiated BPSC induced with TSM for 2 days and ESC. (E) Representative image of ESC and BPSC-diff-D2 chimera at E12.5 stage.

**Figure S4.**
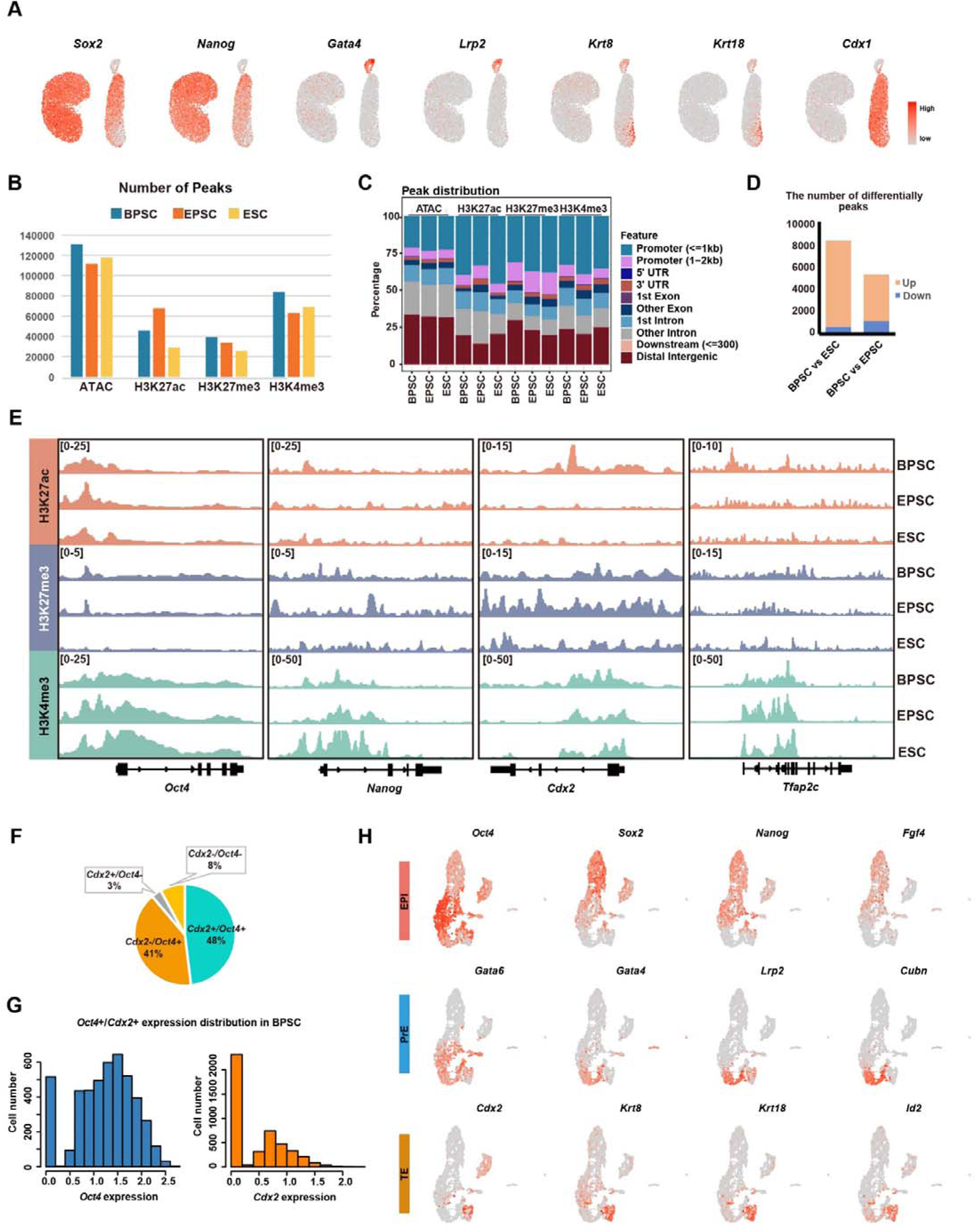
BPSCs exhibit a unique pluripotency state. Related to Figure 3. (A) Gene expression of *Sox2*, *Nanog*, *Gata4*, *Lrp2*, *Krt8*, *Krt18*, and *Cdx1* in BPSC, EPSC, and ESC. (B) Bar plot showing the number of peaks identified from scATAC-seq, H3K27ac ChIP-seq, H3K27me3 ChIP-seq, and H3K4me3 ChIP-seq data in BPSC, EPSC, and ESC. (C) Stacked bar plot showing the distribution of peaks identified from scATAC-seq, H3K27ac ChIP-seq, H3K27me3 ChIP-seq, and H3K4me3 ChIP-seq data on the genome in BPSC, EPSC, and ESC. (D) Stacked bar plot showing the number of upregulated and downregulated differentially accessible peaks identified between BPSC and ESC or between BPSC and EPSC. (E) Genome browser view of H3K27ac, H3K27me3, and H3K4me3 histone modification distribution on *Oct4*, *Nanog*, *Cdx2*, and *Tfap2c* in BPSC, EPSC, and ESC. (F) Pie plot showing the distribution of BPSC grouped based on the expression status of *Oct4* and *Cdx2*. (G) Bar plot showing the *Oct4* or *Cdx2* expression distribution of BPSC. The expression of genes is normalized and log-transformed. (H) Representative gene expression of Epi lineage, TE lineage, and PrE lineage in BPSC, EPSC, ESC, and pre-implantation mouse embryos from zygote to E4.5 blastocyst.

**Figure S5.**
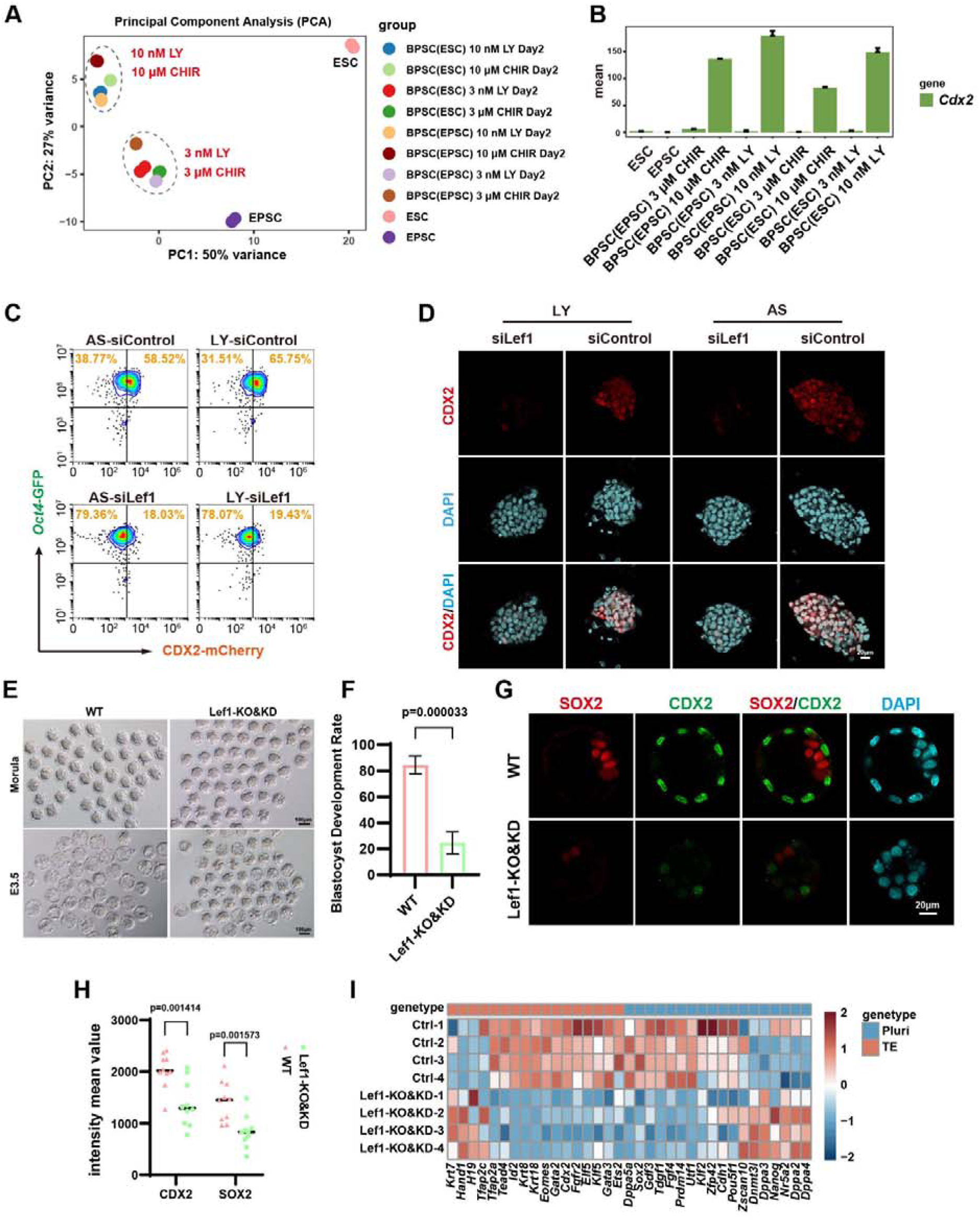
Wnt signalling pathway expands the developmental boundaries of pluripotent stem cells. Related to Figure 4. (A) PCA of bulk RNA-seq data of ESC, EPSC, and ESC or EPSC treated with LY or CHIR for 2 days. (B) Bar plot showing the expression levels of *Cdx2* in ESC, EPSC, and ESC or EPSC treated with LY or CHIR for 2 days. Error bars represent SD. (C) Representative FACS analysis of the percentages of Oct4-GFP positive and CDX2-mCherry positive cells in EPSC treated with AS or LY under knockdown control and knockdown *Lef1* condition. (D) Immunofluorescence staining of EPSC treated with AS or LY under knockdown control and knockdown Lef1 condition. Staining for CDX2. Scale bar, 20 μm. (E) Brightfield images of wild-type (WT) and Lef1 knockout & knockdown morulae and E3.5 embryos. (F) Blastocyst rate of wild-type (WT) and Lef1 knockout & knockdown embryos. (G) Immunofluorescence staining of wild-type (WT) and Lef1 knockout & knockdown E3.5. Staining for CDX2 and SOX2. Scale bar, 20 μm. (H) Statistical analysis of CDX2 and SOX2 fluorescence intensity in wild-type (WT) and Lef1 knockout & knockdown E3.5 embryos. (I) Heatmap showing the expression of TE and pluripotency genes in the bulk RNA-seq of normal blastocyst and *Lef1* knockout/knockdown blastocyst.

**Figure S6.**
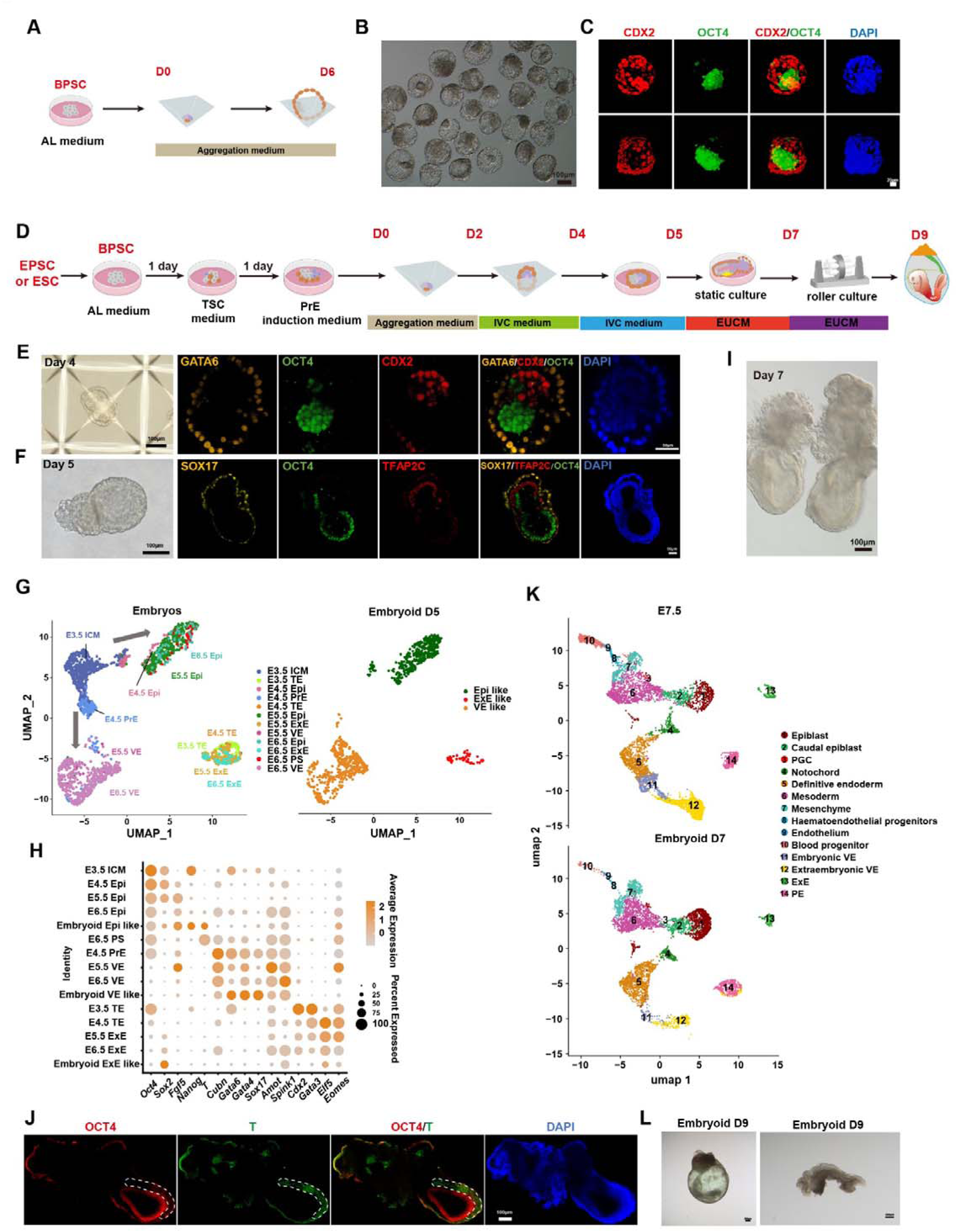
BPSCs alone can form blastoids and early embryonic organogenesis models. Related to Figure 5. (A) Schematic of blastoid construction. (B) The brightfield of blastoids. (C) Immunofluorescence staining of blastoids. Staining for CDX2 and OCT4. Scale bar, 20 μm. (D) Schematic for constructing embryo-like structures (embryoids) with BPSC. (E) Representative morphological images and immunofluorescence staining of embryo-like structures constructed with IVC medium for 4 days. Staining for GATA6, OCT4, and CDX2. Scale bar, 100 μm and 50 μm. (F) Representative morphological images and immunofluorescence staining of embryo-like structures constructed with IVC medium for 5 days. Staining for TFAP2C, OCT4, and SOX17. Scale bar, 100 μm and 50 μm. (G) UMAP of embryoid D5 derived from BPSCs (right panel) and natural E3.5–E6.5 embryos (left panel). (H) Dotplot showing representative gene expression of lineage-specific markers in embryoids and natural embryos. (I) Representative morphological image of E6.5-7.5 like embryoids. Scale bar, 100 μm. (J) Immunofluorescence staining of E7.5 like embryoid. Staining for OCT4 and T. Scale bar, 100 μm. (K) UMAP of natural E7.5 embryos (top panel) and E7.5-like embryoid derived from BPSCs (bottom panel). (L) Brightfield images of embryoid D9.

**Figure S7.**
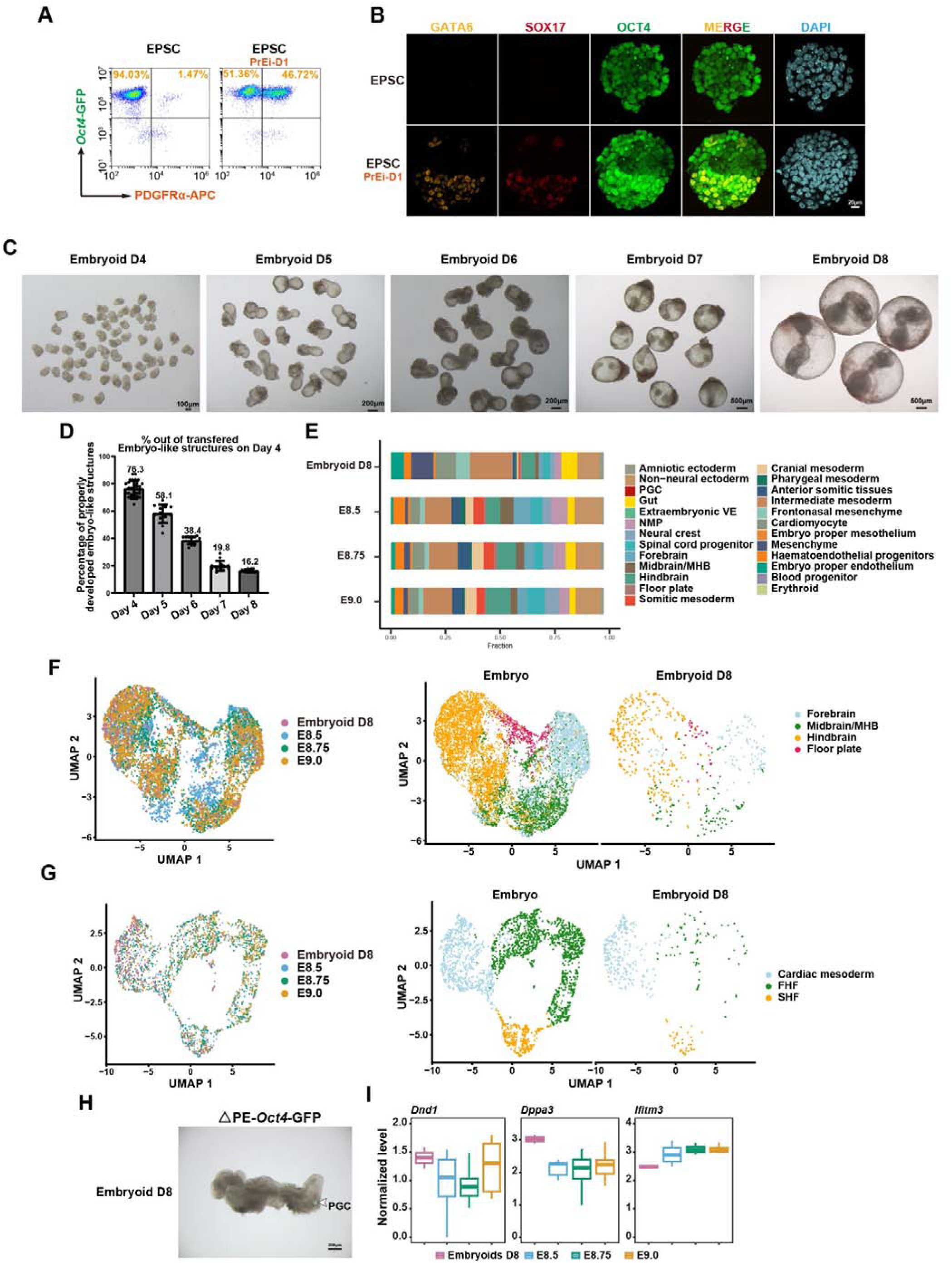
Embryoid D8 shows similar developmental morphology to natural embryos. Related to Figure 5. (A) Representative FACS analysis of the percentages of *Oct4*-GFP and PDGFRα-APC in EPSC and EPSC treated with PrE induction medium one day. (B) Immunofluorescence staining of EPSC and EPSC treated with PrE induction medium one day. Staining for GATA6, SOX17 and OCT4. Scale bar, 20 μm. (C) Bright field images of embryoids developing from Day4 to Day8. (D) Calculated efficiency of properly developed embryoids. The efficiency of well-developed embryoids, relative to the total aggregates on day 4. Error bars represent the SE. (E) Stacked bar plot showing the percentages of different cell types in embryo proper from natural E8.5-E9.0 embryos and embryoids derived from BPSCs. (F) UMAP of brain-related cells in natural E8.5-E9.0 embryos and embryoids derived from BPSCs colored by the stages (left panel) or cell types (right panel). (G) UMAP of heart-related cells in natural E8.5-E9.0 embryos and embryoids derived from BPSCs colored by the stages (left panel) or cell types (right panel). (H) △PE-*Oct4*-GFP fluorescence reveals the location of PGCs in D8 embryoid. (I) Boxplot showing the expression levels of PGC marker genes in PGC from natural E8.5-E9.0 embryos and embryoids derived from BPSCs. The center line represents the median, the box limits indicate the interquartile range.

**Figure S8.**
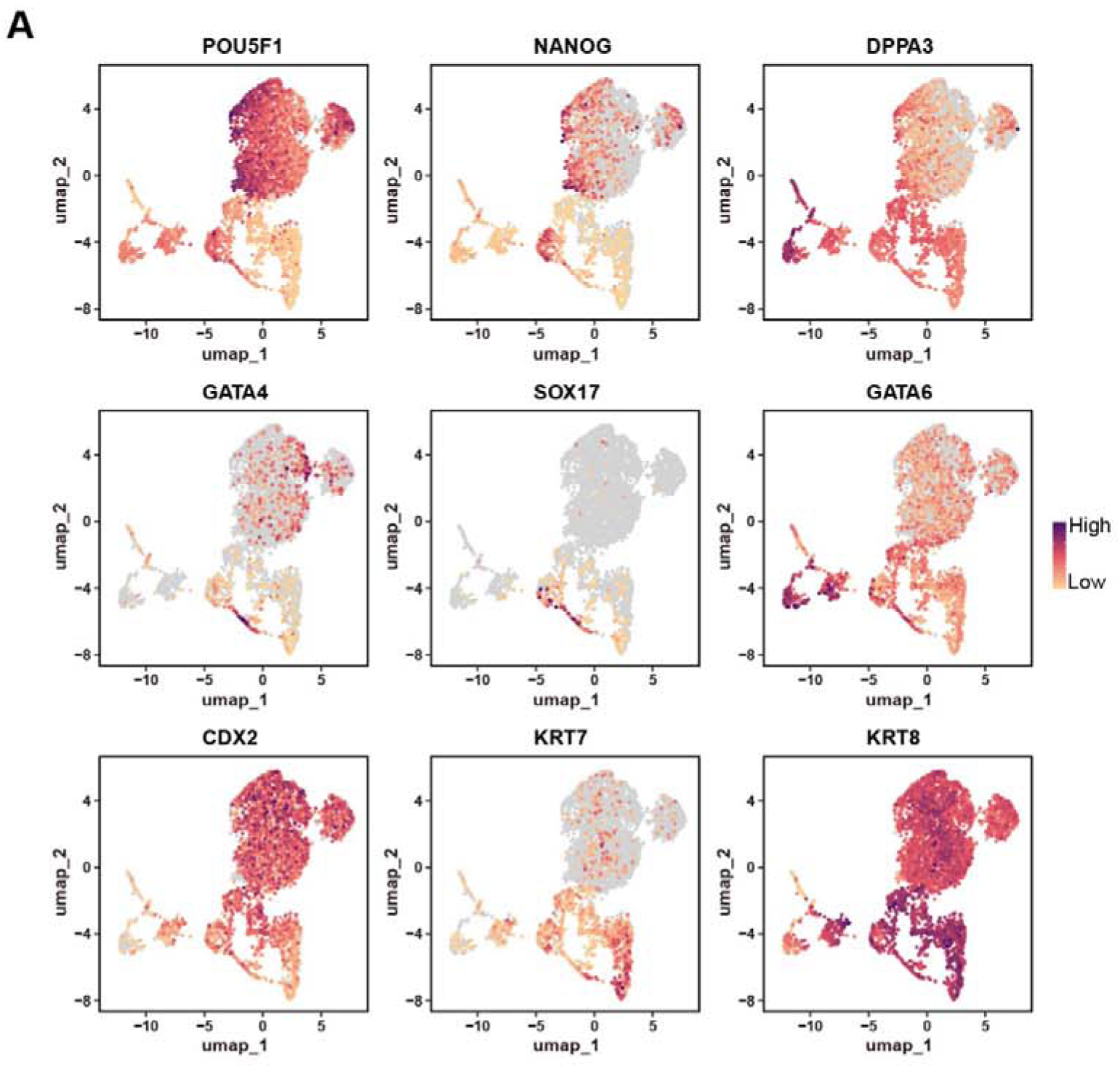
Induction of human BPSCs. Related to Figure 6. (A) Representative gene expression of Epi lineage, TE lineage, and PrE lineage in AL treated nESCs and natural blastocysts.

